# Adaptive Evolution of Gene Regulatory Networks in Mammalian Neocortical Neurons

**DOI:** 10.1101/2025.05.20.652233

**Authors:** Zhuo Li, Navjot Kaur, Gabriel Santpere, Sydney K. Muchnik, Suvimal Kumar Sindhu, Cai Qi, Mikihito Shibata, Olivier Clément, Thomas S. Klarić, Xabier de Martin, Victor Luria, Hyesun Cho, Mingfeng Li, Akemi Shibata, Andrew T. N. Tebbenkamp, Shaojie Ma, Wenqi Han, Suel-Kee Kim, Sirisha Pochareddy, Phan Q. Duy, Xiaojun Xing, Yunhua Bao, Xuming Xu, Ivan Enghian Gladwyn-Ng, Hayley Daniella Cullen, Annalisa Paolino, Laura R. Fenlon, Peter Kozulin, Rodrigo Suárez, Ryan D. Risgaard, Forrest O. Gulden, Amir Karger, Ikuo K. Suzuki, Tatsumi Hirata, Kevin T. Gobeske, Linda J. Richards, André M. M. Sousa, Julian I. Heng, Nenad Sestan

## Abstract

Mammals have evolved a plethora of adaptations that have enabled them to thrive in diverse environments. Among the most significant is the emergence of a more complex brain, exemplified by the dramatic transformation of the dorsal cortex from a single layer of excitatory projection neurons (ExNs) in ancestors to a multilayered cerebral neocortex enriched with diverse intratelencephalic (IT) and extratelencephalic (ET) ExN subtypes. These ExNs established specialized projection systems, such as the corticospinal tract and corpus callosum, enhancing brain connectivity and functionality. However, the evolutionary mechanisms underlying these mammalian-specific adaptations remain elusive. By comparing the landscape of gene expression and cis-regulatory elements (CREs) in mouse ExN subtypes and by cross-species examination of mammalian and non-mammalian CREs, we identified mammalian-specific CREs and expression patterns. The mammalian-specific CREs include a subset bound by ZBTB18 that are associated with genes defining IT and ET subtypes and connectivity. Both ZBTB18 and these target genes have previously been implicated in intellectual disability and autism. Deletion of *Zbtb18* in mouse ExNs dysregulated target gene expression, reduced molecular diversity, diminished corticospinal and callosal projections, and increased intrahemispheric cortico-cortical association projections to the prefrontal cortex, resembling features of non-mammalian dorsal pallium. Interestingly, ZBTB18 binding motifs are highly enriched in callosally projecting IT-biased CREs, where they show higher conservation specifically in mammals. This study uncovers critical components and mammalian-specific evolutionary adaptations within a regulatory node essential for neocortical ExN identity and connectivity, with implications for neurodevelopmental and neuropsychiatric disorders.

## Introduction

The archetypal six-layered cerebral neocortex is a defining feature of the mammalian brain, playing an essential role in advanced cognitive functions ^1–4^. The evolution of the neocortex is characterized by a dramatic increase in both the number ^4^ and the subtype-specific diversity of its principal cells, the excitatory projection neurons (ExNs), also referred to as pyramidal neurons ^5–7^. Additionally, mammalian neocortical ExNs established specialized long-range axonal projection systems, such as the corticospinal tract (CST) and corpus callosum (CC) ^5–8^, enhancing brain connectivity and functionality. In stark contrast, the homologous region of the dorsal pallium in modern reptiles and birds, which are mammals’ closest relatives, is characterized by a single layer or a pseudo-layered columnar organization of predominantly intrahemispheric cortico-cortical and cortico-thalamic projecting ExNs ^4,9–12^.

Previous studies have identified transcription factors (TFs) that guide cell subtype specification, laminar positioning, or connectivity patterns of mammalian neocortical ExNs ^5–7^. However, the evolutionary adaptations and precise molecular mechanisms underlying the defining features of the mammalian neocortical ExNs remain elusive. Here, we conducted transcriptome sequencing (RNA-Seq) and chromatin immunoprecipitation followed by sequencing (ChIP-Seq) on genetically labeled populations of two major classes of developing mouse neocortical ExNs, the intratelencephalic-projecting (IT) and the extratelencephalic-projecting (ET) neurons, and compared this to gene expression and regulatory element profiling in chicken dorsal pallium. This uncovered both common and subtype-specific *cis*-regulatory elements (CREs) in developing ExNs that confer uniquely mammalian attributes. This included a notable subset of CREs bound by ZBTB18 (also known as RP58, ZFP238, or ZNF238) that are exclusively found in mammals or exhibit either unique mammalian characteristics and are associated with TFs defining ET/IT neuron subtypes. Prior research has established that ZBTB18 is enriched in the developing mammalian neocortex and plays a pivotal role in regulating neurogenesis and neuronal migration ^13–17^, while mutations in the human *ZBTB18* gene are linked to intellectual disability and autism ^18–22^. This prompted further exploration of ZBTB18’s postmitotic functions in the neocortical ExNs of mice and investigation of conservation rates of binding motifs within ET/IT-biased CREs that may indicate unique evolution in mammals. These experiments collectively uncovered that gene regulatory subcircuits, particularly involving ZBTB18-CRE interactions, which govern key features of the identity and connectivity of neocortical ExNs, have undergone modifications in the mammalian lineage.

## Results

### Subtype-specific CREs and TFs in mouse neocortical ExNs

To characterize CREs and TFs shaping the diversification of neocortical ExNs, we utilized *Arpp21-Gfp or Fezf2-Gfp* transgenic mice having GFP-expressing neurons enriched in the neocortical upper layer (L2-4) IT neurons or deep layer (L5-6) predominantly ET neurons, respectively (**Fig. 1a**). GFP-expressing cells were collected using fluorescence-activated cell sorting (FACS) from neonatal mice (postnatal day 0, PD 0 hereafter), an age when neocortical ExN identity and connectivity were being established. Collected cells were then processed for RNA-Seq and ChIP-Seq using an antibody against H3K27ac, a marker of active enhancers and promoters^23–27^ (**Fig. 1b**).

**Figure 1.**
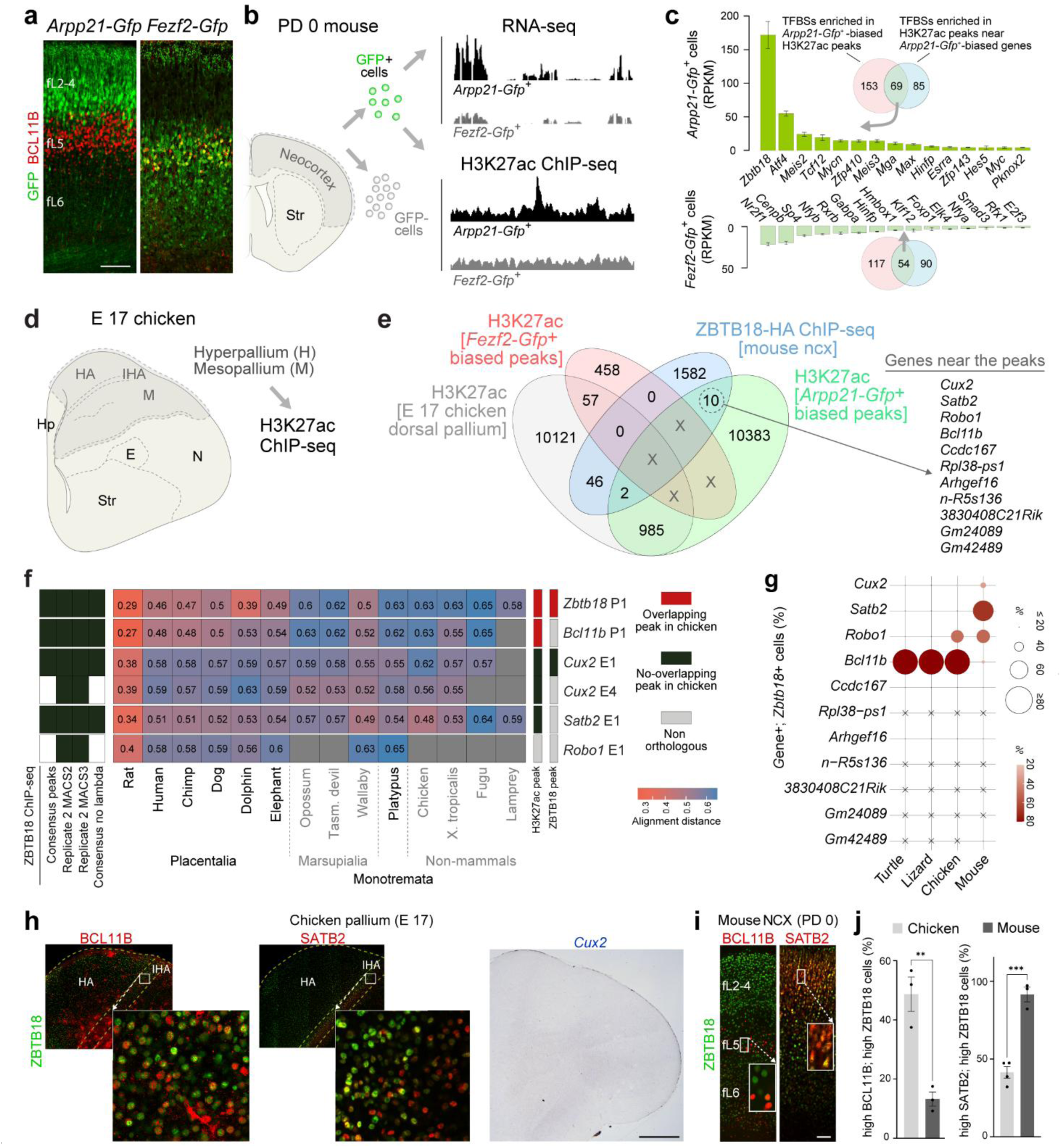
Mammalian-specific changes in the ZBTB18-associated CREs and TF expression in neocortical ExNs. **a**) The expression of GFP (green) driven by *Arpp21* and *Fezf2* CREs and BCL11B immunolabeling (red) in mouse PD 0 neocortex. fL, fetal (immature) layer. Scale bar: 100µm. **b**) Schematic depicting the isolation of GFP-labeled cells from *Arpp21-Gfp* and *Fezf2-Gfp* neocortex and processing for RNA-Seq and H3K27ac ChIP-Seq. **c**) TFBS for 69 TFs were enriched among both *Arpp21-Gfp*-biased H3K27ac peaks (red circle) and H3K27ac peaks near *Arpp21-Gfp*-biased genes (blue circle) as compared to ET neuron-biased peaks and genes, respectively. TFBS enrichments were tested using Fisher’s exact test. Significant TF motifs had a Benjamini-Hochberg corrected p-value < 0.05 and odds ratio > 1. Among these 69 TFs, *Zbtb18* was the most highly expressed in IT neurons. For RNA-Seq, a minimum of 3 independent biological replicates were analyzed for each condition. For ChIP-Seq, 2 biological replicates were analyzed for each condition. **d**) Schematic showing two dorsal pallial regions (H and, M) microdissected from embryonic day (E) 17 chicken embryo. Hp (hippocampus), H (hyperpallium), HA (apical hyperpallium), IHA (interstitial apical hyperpallium), M (mesopallium), N (nidopallium), E (entopallium), Str (striatum). **e**) Venn diagram showing the co-occurrence of peaks derived from ZBTB18-HA ChIP-Seq, and H3K27ac peaks preferably found in *Arpp21-Gfp*+ IT neurons, preferably in *Fezf2-Gfp*+, and in chicken dorsal pallium. 10 peaks appear in the ZBTB18-HA ChIP-Seq samples that were exclusively present in the *Arpp21-Gfp*+ IT neurons. To meet the IT or ET preference criteria, peaks should be present in both samples of one group and a maximum of one sample from the other group. X represents the common peaks that were present in both *Arpp21-Gfp*+ IT neurons and *Fezf2-Gfp*+ ET neurons and were not included in the analysis. **f**) Heatmap illustrates the pairwise alignment distances between various species of vertebrates for the six putative CREs that overlap with ZBTB18 ChIP-Seq peaks and are associated with IT-neurons and axon guidance molecules. The ChIP-Seq analysis method employed to identify each peak is indicated in the left grid (see methods). Additionally, columns on the right provide information on whether the corresponding region (H3K27ac peak - left column or ZBTB18 ChIP-seq peak - right column) exhibits orthology and H3K27ac activity in the chicken embryonic dorsal pallium. The grey columns depict the non-orthologous regions between the mouse and the chicken. The red and black column depicts the presence or absence of an overlapping H3K27ac peak in the regions that are orthologous in mice and chicken, respectively. Alignment distance of various species of vertebrates is measured from mice. **g**) Dot plot showing the percentage of cells among *Zbtb18* expressing neurons with the dorsal pallium that co-express the genes carrying ZBTB18 binding motif within their loci in *Arpp21-Gfp*+ neurons amongst mammalian (mouse) and non-mammalian species (chicken, lizard, and turtle) ^12,29–31^. **h-i**) Coronal sections of chicken and mouse brain showing the extent of colocalization of BCL11B and SATB2 with ZBTB18. Scale bar: 1mm (mouse brain), 150 µm (mouse inset), 1mm (chicken brain), 300 µm (chicken inset), 250 µm (chicken brain; *Cux2*). (j) Bar plots showing the percentage of the co-localization of BCL11B and ZBTB18 over total BCL11B-immunopositive cells and SATB2 and ZBTB18 over total SATB2-immunopositive cells in chicken and mouse. A standard t-test, unpaired, was applied. The graph represents mean ± s.e.m. (standard error mean). ** P = 0.0011, *** P = 0.0004 (n = 3/ species).

Commensurate with the shared developmental lineage of IT and ET neurons, ChIP-Seq showed that the majority of the expressed genes [15,120 out of 15,902] and the genomic regions enriched for H3K27ac sites (peaks) [35,173 out of 62,448] were common to both presumptive *Arpp21-Gfp*+ IT neurons and *Fezf2-Gfp*+ ET neurons (**Extended Data Table 1-2**). Gene ontology analysis of the genes and ChIP-Seq peaks enriched specifically in either *Arpp21-Gfp*+ cells [304 genes and 54 IT neuron-enriched peaks] or *Fezf2-Gfp*+ cells [478 genes and 73 ET neuron-enriched peaks] (**Extended Data Fig. 1a-c, Extended Data Tables 1-3**) revealed terms broadly relevant for either IT neurons or ET neurons functions, respectively (**Extended Data Fig. 1d, Extended Data Tables 4-7**).

To prioritize TFs associated with the development of IT and ET neurons, we searched for those that are highly and differentially expressed and whose TF binding sites (TFBS) are enriched among H3K27ac peaks specific to *Arpp21-Gfp*+ cells (222 TFs, **Fig. 1c, Extended Data Table 8**). We also highlighted TFBS that are proximal to genes upregulated in the same cells (154 TFs), compared to *Fezf2-Gfp+* cells, and vice versa (**Fig. 1c, Extended Data Table 9**). Some of the prioritized TFs have been previously implicated in the development of IT and ET neurons ^5–7^, including *Meis2, Rara, Rarb*, and *Rxrg*, which mediate the retinoic acid signaling during the prefrontal cortex (PFC) development^28^. In *Arpp21-Gfp*+ cells, 69 TFs had binding motifs enriched in ChIP-Seq datasets and also had enhanced expression in RNA-seq datasets. Amongst these, ZBTB18 was the most highly expressed TF in *Arpp21-Gfp*+ cells (**Fig. 1c, Extended Data Table 8**). Similarly, 54 TFs had binding motifs enriched and increased expression in *Fezf2-Gfp+* cells, with NR2F1 being the most highly expressed in them (**Fig. 1c, Extended Data Table 9**). Through our integrated analysis, we discovered both shared and cell type-specific TFs and CREs. Notably, our findings highlighted ZBTB18, prompting us to investigate its regulatory network and functions in developing neocortical ExNs.

## ZBTB18 targets mammalian-specific enhancers of key ExN genes

Upon identifying ZBTB18 as a TF with potentially important roles in ExN sub-specification, we sought to validate the regulatory sites in the developing mouse neocortex occupied by ZBTB18 using ChIP-Seq analysis. Yet because our initial trials with several commercially available anti-ZBTB18 antibodies could not yield suitably high-quality and reproducible ChIP-Seq findings, despite working well for immunostainings (**Extended Data Fig. 4a**), we conducted ChIP-Seq by expressing HA-tagged mouse *Zbtb18* plasmid electroporated into PD 0 neocortical ExNs. This analysis demonstrated that ZBTB18 binds to developmental CREs marked by H3K27ac peaks near crucial genes previously implicated in the development and diversification of ExNs, such as *Bcl11b*, *Cux2,* and *Satb2* (**Extended Data Table 11**).

To explore potential roles that ZBTB18-based gene regulation may play in the mammalian evolution of the neocortex, we compared the mouse ZBTB18 ChIP-Seq dataset and the H3K27ac peaks from *Arpp21-Gfp+* and *Fezf2-Gfp+* cells with H3K27ac peaks derived from a non-mammalian species, the chicken (**Fig. 1d, Extended Data Table 10**). This analysis was performed on micro-dissected hyperpallium (H), and mesopallium (M), which are regions within the avian dorsal pallium considered to be homologous to the mammalian neocortex, based on multiple lines of evidence, including gene expression profiles ^4,10,11^. We isolated these regions from chicken embryos at embryonic day (E) 17, corresponding approximately to a developmental age equivalent to that of a PD 0 mouse. By conducting phylogenetic analysis and comparing the ChIP-seq datasets - including H3K27ac from *Arpp21-Gfp+* IT neurons, *Fezf2-Gfp+* ET neurons, and ZBTB18-HA from mouse neocortical neurons, as well as H3K27ac from the chicken dorsal pallium -we identified 10 distinct enhancers or promoters wherein ZBTB18 binds that were also biased to *Arpp21-Gfp+* IT neurons of mice but not in the chicken dorsal pallium (**Fig. 1e, Extended Data Table 11**). Among the 10 ZBTB18-bound peaks that exhibited exclusive accessibility in mice but not chickens, 5 were identified in proximity to crucial genes differentially expressed between IT and ET neurons and previously implicated in their development ^5–7^: *Cux2* (referred to as *Cux2* enhancer (E) E1 and *Cux2* E4), *Satb2* (*Satb2* E1), *Robo1* (*Robo1* E1), and *Bcl11b* (*Bcl11b* promoter (P) P1) (**Fig. 1e-f, Extended Data Fig. 2a-c, Extended Data Fig. 9a**). To assess the conservation of these ZBTB18-bound peaks, we employed sequence identity measurements extracted from MULTIZ60 multiple species alignments, along with H3K27ac peaks derived from the embryonic chicken dorsal pallium. Remarkably, within the ZBTB18-bound peak associated with mouse *Cux2* E1, no overlapping H3K27ac peaks were detected in the orthologous region in chickens. This observation suggests that this non-coding region serves as an active enhancer in the mouse cortex but not in the chicken (**Fig. 1f, Extended Data Fig. 2a**). Regarding the ZBTB18-bound peaks within mouse *Cux2* E4 and *Satb2* E1, we were unable to identify orthologous regions or overlapping H3K27ac peaks in the chicken homologous region (**Fig. 1f, Extended Data Fig. 2a-b**). As a result, we can exclude them as enhancers in the E17 chicken dorsal pallium.

Conversely, the ZBTB18-bound peak within the mouse *Bcl11b* P1 did not have an orthologous region in chickens (**Fig. 1f, Extended Data Fig. 2c**). However, the surrounding regions did show the signal in H3K27ac ChIP-seq for chicken dorsal pallium, suggesting that it functions as an active promoter in chickens, and this activity is not potentially regulated by ZBTB18. In contrast, we could not identify an orthologous region for mouse *Robo1* E1 in chickens or any other non-mammalian species we analyzed (**Fig. 1f, Extended Data Fig. 9a**). This indicates that non-mammalian species lack this putative enhancer. Interestingly, we also identified ZBTB18-bound and H3K27ac peaks within corresponding regions in mouse ExNs and the chicken dorsal pallium, including the promoter region of *Zbtb18* (**Extended Data Fig. 2d**). This observation hints at a potential evolutionarily conserved autoregulatory role for ZBTB18. These findings also collectively indicate that specific ZBTB18-bound neocortical ExN subtype-biased enhancers, which are associated with key regulators of ExN development, exhibit distinct mammalian features. This suggests that ZBTB18 may play a role in the evolution of neocortical ExN diversification in mammals.

For further review of ZBTB18-interacting regions we selected *Cux2* E1 (1,036 bp in size) and *Satb2* E1 (4,474 bp in size) that contain consensus binding sites for ZBTB18 in all analyzed mammals, and which, in the chicken, either lacked H3K27ac peaks or the regions associated with the ZBTB18-bound peak was absent in the genome, respectively (**Extended Data Fig. 2a-b**). In humans, the putative enhancers identified for *CUX2* and *SATB2* exhibited H3K27ac enrichment in fetal dorsolateral PFC(dlPFC or DFC) ^25^, suggesting that the activation of these enhancers is conserved in the mammalian developing neocortex (**Extended Data Fig. 2a-b**). Additionally, we analyzed the co-expression pattern of *Zbtb18* with genes that have ZBTB18-bound peaks in *Arpp21-Gfp*+ IT neurons (**Fig. 1e**) using publicly available, single-cell RNA-seq datasets for the mouse cortex and the dorsal pallium of birds, lizards, and turtles ^12,29–31^. Among these genes, *Zbtb18* exhibited the highest co-expression with *Cux2*, *Satb2,* and *Robo1* in mice, but not in non-mammalian species (**Fig. 1g**). Conversely, in chickens, lizards, and turtles, *Zbtb18* showed the highest co-expression with *Bcl11b*, an ET-specific TF in mammals, suggesting a stronger association with IT neurons in mammals but potentially less ExN class selectivity earlier in evolution (**Fig. 1g)**.

Further, using anti-ZBTB18 antibodies that were independently validated (**Extended Data Fig. 4a**), we conducted immunolabeling on ExNs in mouse, and we observed that ZBTB18 exhibited strong co-localization with IT-enriched TFs, SATB2 and *Cux2*, but not with BCL11B **(Fig. 1i-j, Extended Data Fig. 4b)**. Immunohistochemical analysis of the embryonic chicken pallium revealed regionally extensive ZBTB18 expression, with a higher density of labeled nuclei in the HA (apical hyperpallium), particularly in the interstitial apical hyperpallium (IHA), the main sensory input subregion of the HA, which contains neuronal types molecularly resembling mammalian neocortical thalamorecipient IT ExNs ^11^ (**Fig. 1h, Extended Data Fig. 1e**). Consistent with previous gene expression analyses in the embryonic chick pallium ^4,11^, we observed expression of BCL11B and *Fezf2* that were stronger in the HA region than in the M region, whereas SATB2 was expressed predominantly in the M region compared to HA (**Extended Data Fig. 1e**). In contrast to the co-expression pattern observed in the developing mouse neocortex, ZBTB18 exhibited increased co-localization with BCL11B rather than SATB2 in the HA region, IHA region and adjacent M region in the chicken pallium (**Fig. 1h-j, Extended Data Fig. 1e**). Surprisingly, we did not detect appreciable *Cux2* expression in most regions of the dorsal pallium, except in the posterior ventral pallium region (**Fig. 1h, Extended Data Fig. 1e**). Our analysis of single-cell RNA-seq datasets ^12,29–31^ also revealed little or no appreciable *Cux2* expression in *Zbtb18*-expressing neurons within the dorsal pallium of birds, lizards, and turtles, unlike the developing mouse neocortex where the *Cux2* and *Zbtb18* are highly co-expressed (**Fig. 1g**). Therefore, in mammals, ZBTB18 may regulate the expression of genes such as *Cux2, Satb2,* and *Robo1* by binding to specific sites within the *Cux2* E1, *Satb2* E1 and *Robo1* E1 enhancers, which are not active enhancers or are absent in non-mammalian species. From an evolutionary perspective, this suggests that ZBTB18 plays a crucial role in mediating regulatory networks contributing to the enhanced diversity and connectivity of neocortical ExNs observed in the mammalian lineage.

### ZBTB18 regulates mammalian *Cux2* neocortical enhancer

Because *Cux2* expression is specific to IT neurons in mice, with only minimal expression in the embryonic chicken dorsal pallium (**Fig. 1h**, **Extended Data Fig. 1e**) and given that the H3K27ac peaks in *Arpp21-Gfp+* IT neurons were prominent for *Cux2* (**Extended Data Fig. 2a**), we selected *Cux2* as a potential target for ZBTB18 during the specification of IT neurons. To validate the activity of this enhancer, we generated multiple transgenic founders in which *Cux2* E1 was placed 5’ to the human *BGN* promoter and was linked with a *Gfp* reporter gene. In these mice, we observed GFP expression in the forebrain by PCD 14.5, recapitulating the native expression of *Cux2* ^32,33^ (**Extended Data Fig. 3a**). By PCD 16.5, forebrain GFP expression was restricted to a subset of neurons in the neocortical plate (**Fig. 2a-b**), with such expression co-localizing with the ZBTB18 native protein in the neocortex but not the striatum (**Fig. 2b**). Consistent with the importance of *Cux2* E1 for gene expression broadly in the neocortex and specifically in IT neurons, we found that the expression of *Cux2* E1*-Gfp* also co-localized with the IT neuron marker protein SATB2, but not the ET neuron marker BCL11B at PCD 16.5, 17.5 and PD 0 (**Fig. 2c-d, Extended Data Fig. 3b-c)**. By PD 15, most GFP labeled cells were located in L2 through L4, and were immunopositive for CUX1, a marker frequently co-expressed by CUX2-expressing cells, but were not immunopositive for BCL11B (**Extended Data Fig. 3d-f**). Also matching these findings, we observed GFP in the CC but not in the CST, which is a major projection system of L5 ET neurons, when it was surveyed at the pontine region on the ventral surface of the brain at PD 0 (**Extended Data Fig. 3c**). In contrast to the specificity of *Cux2 E1*, two putative *Cux2* enhancers, *Cux2* E2 and *Cux2* E3 were active in both IT and ET neurons and lacked ZBTB18 ChIP-seq peak, hence subsequently being used as negative controls (**Extended Data Fig. 3g-h**). Also as expected, *Cux2* E2 and *Cux2* E3 were unable to drive expression selectively and strongly in IT neurons, including callosally projecting subsets, but instead had expression mainly in RELN-immunopositive L1 neurons and LHX6-immunopositive interneurons, respectively (**Extended Data Fig. 3g-h**).

**Figure 2.**
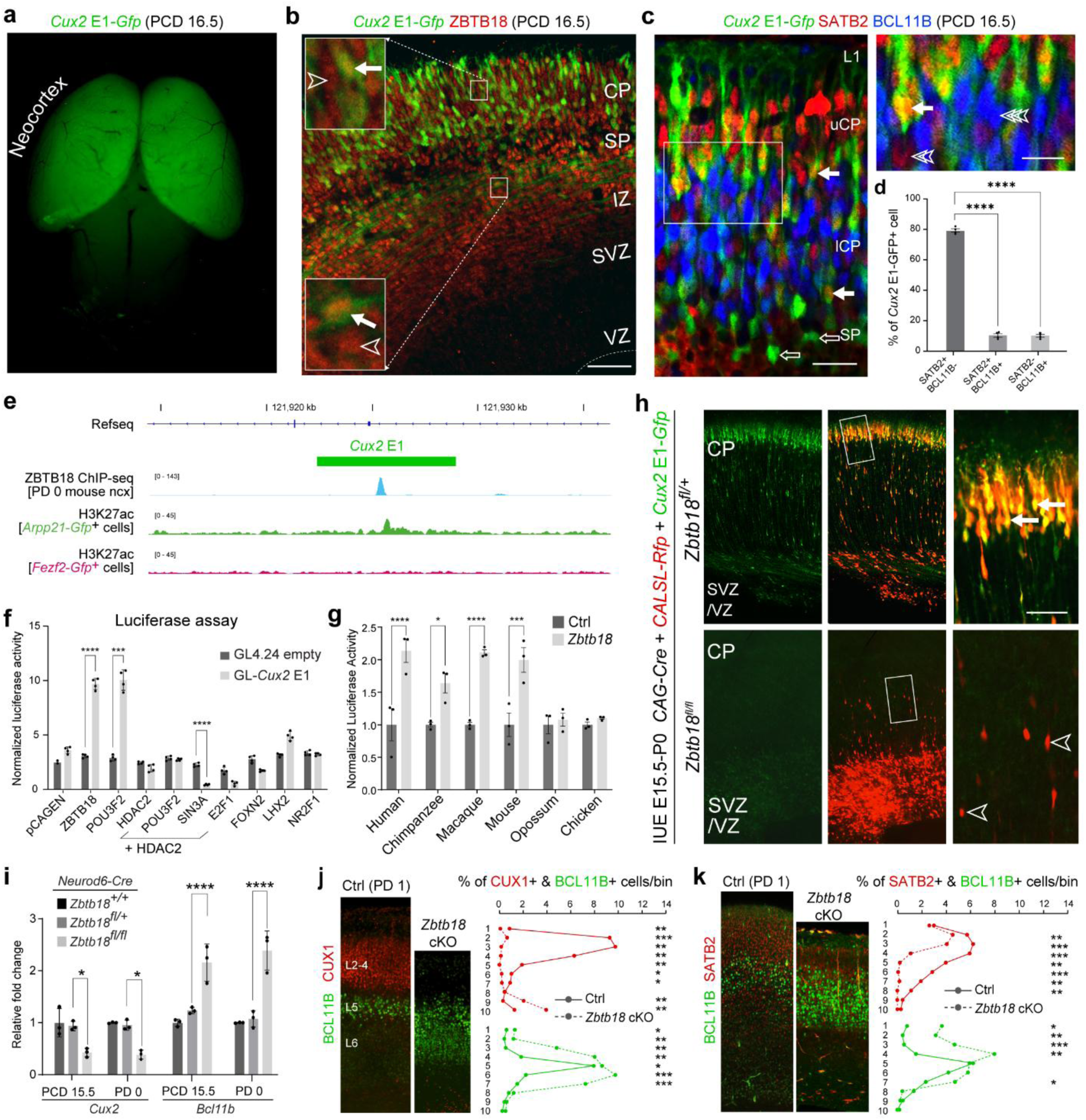
ZBTB18 directly regulates mammalian-specific neocortical enhancer of *Cux2*. **a**) *Cux2* E1 drives *Gfp* expression in the PCD 16.5 neocortex. Scale bar: 1mm. **b**) *Cux2* E1*-Gfp* expression co-localizes with ZBTB18 (closed arrows) in the neocortex. Several ZBTB18+ cells do not express GFP (open arrowheads). Scale bar: 100 µm. **c-d**) At PCD 16.5, *Cux2* E1 mediated expression of GFP overlaps primarily with SATB2 (closed arrows) but not BCL11B. Double open arrowheads (SATB2) and triple open arrowheads (BCL11B) indicate cells immunolabeled for these markers but not *Cux2* E1-*Gfp*. Ordinary one-way ANOVA with Dunnett’s multiple comparisons, with a single pooled variance was used. An unpaired, two-tailed t-test was used to detect differences between groups (d). The graph represents mean ± s.e.m. (standard error mean). * P= 0.0001. Scale bars: 200µm; Inset, 50 µm. **e**) Line graphs showing the H3K27ac peaks from *Arpp2-Gfp*+ IT neurons, *Fezf2-Gfp*-positive ET neurons and ZBTB18-HA ChIP-Seq peaks associated with mouse *Cux2*. **f**) Luciferase activity driven by the *Cux2* E1 enhancer is significantly increased by ZBTB18. An unpaired t-test was used to detect differences between control and experimental conditions. The graph represents mean ± s.e.m. * P = 0.000040, 0.00029, 0.000057 (ZBTB18, POU3F2, HDAC2+ SIN3A). **g**) Luciferase activity driven by the *Cux2* E1 enhancer from different species is significantly increased by ZBTB18 only in placental mammals (Eutherians). Ordinary two-way ANOVA with Bonferroni’s multiple comparisons test. The graph represents mean ± s.e.m. **** P < 0.0001, ***= 0.002, *= 0.168 (n = 3). **h**) *Cux2* E1 driven expression of GFP is lost following the conditional deletion of *Zbtb18*; although *Cux2* E1*-Gfp* co-localizes with electroporated, RFP+ cells in control *Zbtb18* ^fl/+^ brain (closed arrows), many cells where *Zbtb18* has been deleted (red) fail to express GFP and remain in the proliferative niche (open arrowheads), evidencing a proliferation or migration defect. Scale bar: 50 µm. **i**) Conditional deletion of *Zbtb18* in ExNs resulted in a significant downregulation of *Cux2* and the upregulation of *Bcl11b* at both PCD 14.5 and PD 0. Ordinary two-way ANOVA with Bonferroni’s multiple comparisons test, with single pooled variance, was applied. The graph represents mean ± s.e.m. * P= 0.0327, 0.0178 (WT vs *Zbtb18* cKO at E15.5, and PD 0), P = 0.0001 (WT vs *Zbtb18* cKO at E15.5, and PD 0). The native distribution of CUX1 or SATB2, respectively, (red, upper layers) and BCL11B (green, deep layers) is disrupted by the conditional deletion of *Zbtb18* in *Neurod6-Cre*; *Zbtb18* cKO brains at PD 1. Scale bar: 100 µm. **j-k**) Quantification of cell-type laminar distribution from b and c, respectively. T-tests were used to compare control and knockout cell counts per bin. mean ± s.e.m at each bin. For the p-value, see Supplementary Table 1. For RNA-Seq and RT-PCR n = 3/ timepoint. For immunofluorescent analyses, we counted neurons from independent sections (n = 3/ condition).

We next conducted luciferase activity-based enhancer assays to determine which TFs might be responsible for the observed neocortical *Cux2* E1 activity. We identified several TFs, including ZBTB18, that are expressed in the developing mouse and human forebrain and exhibit binding motifs within these regions, as predicted by multiple algorithms. Of the tested TFs, ZBTB18 and POU3F2/BRN2 increased *Cux2* E1 activity, and a combination of HDAC2 and SIN3A repressed the activity (**Fig. 2f**). Combined with findings from the ChIP-seq (**Fig. 2e)** demonstrating that ZBTB18 directly binds to *Cux2* E1, these results strongly suggest that ZTBT18 acts as a direct activator of *Cux2* transcription, at least under *in vitro* conditions. To assess whether *Cux2* E1 enhancer activity is specific to placental mammals, we generated luciferase constructs containing orthologous *Cux2* E1 sequences from human, chimpanzee, macaque, mouse, opossum, and chicken. Notably, only the sequences from placental mammals, but not those from opossum or chicken, exhibited a significant increase in luciferase activity upon ZBTB18 co-expression (**Fig. 2g**), indicating that ZBTB18-dependent *Cux2* E1 enhancer activation is an evolutionary adaptation specific to placental mammals.

We next sought to determine whether ZBTB18 was necessary for the transactivation of *Cux2 E1 in vivo*. For this, we co-electroporated a *Cux2* E1*-Gfp* plasmid, a *CALSL-Rfp* reporter plasmid, and a *CAG-Cre* plasmid into mice carrying homozygous (*fl/fl*) or heterozygous (fl/+) floxed *Zbtb18* alleles. To target IT neurons predominantly, electroporation was performed at PCD 15.5, when only upper-layer IT neurons (mainly callosally projection neurons) are generated. At PD 0, we observed co-localization of GFP and RFP in upper layer neurons in the *Zbtb18^fl/+^* brain. By contrast, we observed RFP without commensurate GFP expression in electroporated *Zbtb18*^fl/fl^ cells (**Fig. 2h**), indicating ZBTB18 is required for *Cux2* E1 transactivation.

In addition to *Cux2* E1, we also identified a consensus binding site for ZBTB18 within *Satb2* E1 (**Extended Data Fig. 2b**). Thus, to test the specificity of the ZBTB18 binding site within *Satb2* E1 CRE, we conducted luciferase assays by co-transfection of the *Zbtb18* expression construct with DNA constructs for *Satb2* E1, or *Satb2* ΔE1(mutant construct with no ZBTB18 binding site). This demonstrated that *Zbtb18* significantly increased the activity of *Satb2 E1* compared to their respective controls. Although the luciferase reporter activities from the Satb2 ΔE1 constructs were reduced, they did not return entirely to the basal levels (**Extended Data Fig. 2e**).

To assess the extent of conservation in the developing mammalian neocortex, we analyzed putative CREs overlapping with the *Cux2* E1, *Satb2* E1, *Bcl11b* P1, *Zbtb18* P1, and *Robo1* E1 loci in an independent H3K27ac ChIP-Seq database from the human midfetal dlPFC ^25^. Our analysis revealed overlaps between CREs associated with the loci in both human and developing mammalian neocortex (**Extended Data Fig. 2a-d, Extended Data Fig. 9a**). These findings indicate conservation of these CREs in various mammals, suggesting their potential involvement in fundamental neocortical developmental processes. Collectively, these results also highlight the pivotal role of ZBTB18 in regulating such CREs, underscoring the necessity for further comprehensive investigation.

### Enrichment of ZBTB18 in postmigratory IT neurons

Previous studies have shown that the postmigratory stage is critical for ExN specification ^5–7^. To determine the biological relevance of the ZBTB18 functions in the ExNs, we validated its co-expression with cortical ExN markers. Aligning with its function in the direct regulation of *Cux2* E1, ZBTB18 was found to be co-expressed with CUX2 in upper layer IT neurons in the mouse at PD 0 and the human at post-conception week (PCW) 20, which corresponds to the late mid-fetal stage of human development and is equivalent to PD 0 in mice (**Fig. 1i-j, Extended Data Fig. 4b-c**). Furthermore, our analysis of a publicly available mouse neocortical single-cell RNA-seq dataset ^34^ revealed that *Zbtb18* expression was higher in developing IT neurons compared to that in L5 and L6 ET neurons (**Extended Data Fig. 5a**). This expression pattern was replicated in the late mid-fetal human neocortex, where strong immunolabeling for ZBTB18 was found in nuclei positive for SATB2 and CUX2 within upper layer prospective IT neurons (**Extended Data Fig. 4c, Extended Data Fig. 5b-c**). However, the ZBTB18 immunolabelling was diminished in BCL11B-immunopositive nuclei of prospective L5B ET neurons (**Fig. 1i-j, Extended Data Fig. 5b-c**). Therefore, the enrichment and sustained high levels of ZBTB18 expression in IT neurons suggest its crucial role in their specification and differentiation, especially after these cells have completed their postmitotic and migratory phases.

### Postmitotic requirement of ZBTB18 for ExN sub-specification

To investigate the effects of *Zbtb18* on ExN specification in more detail, we next tested the expression of genes enriched in IT neurons vs ET neurons following *Zbtb18* knockout interventions. We performed RNA-Seq on *Zbtb18*^-/-^ (KO) and *Zbtb18*^+/-^ (Control) neocortex at PCD 14.5, a time coinciding with the generation of upper layer callosally projection neurons (**Extended Data Fig. 6a, Extended Data Table 12**). Of the genes that were enriched in either IT or ET neurons and differentially regulated in the *Zbtb18* KO at PCD 14.5, we found that disproportionally more genes enriched in IT neurons were downregulated following knockout (82.3% [90 of 113]) in the *Zbtb18* KO, as opposed to the upregulated genes). Yet the opposite trend was observed with ET neuron enriched genes in the *Zbtb18* KO (92.4% [219 of 237] of which were upregulated) (**Extended Data Fig. 6a, Extended Data Table 12**).To test whether ZBTB18 is required for neuron specification post-mitotically, we conditionally inactivated *Zbtb18* in postmitotic neurons by crossing *Zbtb18^fl/fl^* mice to *Neurod6-Cre* (aka *Nex1-Cre*) transgenic mice that express *Cre* strongly in cells committed to cortical glutamatergic fates; and we then conducted RNA-Seq on neocortical tissue derived from *Neurod6-Cre*; *Zbtb18^fl/fl^* (*Zbtb18* cKO) and *Neurod6-Cre; Zbtb18^fl/+^* (control) mice at PD 0 (**Extended Data Fig. 6b, Extended Data Table 13**). Similar to the PCD 14.5, at PD 0 when all immature layers are formed, 58.7% of genes (47 out of 80) enriched in IT neurons and differentially expressed in the *Zbtb18* KO at PD 0 were downregulated, whereas 96.7% of genes (175 of 181) enriched in ET neurons were upregulated (**Extended Data Fig. 6b, Extended Data Table 13**). *In situ* hybridization, immunohistochemistry, and quantitative PCR (qPCR) analyses confirmed a reduction in the expression of key IT neuron marker genes, such as *Cux1*, *Cux2*, *Rorb*, and *Satb2*, within the neocortical plate of *Zbtb18* KO and cKO mice (**Fig. 2i, Extended Data Fig. 6c, Extended Data Fig. 6h**). Conversely, expression of the ET neuron molecular marker BCL11B was increased in these same mice (**Fig. 2j-k**).

Anatomical and histological analysis of *Neurod6-Cre*; *Zbtb18* cKO mice at PD 0 revealed a minor reduction in neocortical size, which contrasts with findings in the constitutive (whole-body) or *Emx1-Cre* (progenitor and postmitotic levels) KO mice ^13,16^, as compared to control littermates (**Extended Data Fig. 6d**). However, at later postnatal age PD 8, we did observe a substantially smaller neocortical size in the conditional KO mice (**Extended Data Fig. 6d**). Nevertheless, ZBTB18 protein expression in the *Neurod6-Cre; Zbtb18* cKO mice was present within neocortical progenitor cells but absent from post-mitotic cells with the intermediate zone (IZ), subplate (SP) and cortical plate (CP), when analyzed at PCD 15.5 and onward (**Extended Data Fig. 6e**). This observation implies that the effects of *Zbtb18* knockout in this system do not stem from neurogenesis or progenitor-related processes. Post-mitotic actions of *Zbtb18* knockout still can affect other cell populations, though, as we found that levels of CUX2 protein and mRNA in the cortical plate were reduced in *Neurod6-Cre; Zbtb18* cKO neocortex (**Extended Data Fig. 6f-h**) as well as in *Emx1-Cre*; *Zbtb18* cKO mice and whole-body KO when analyzed by in situ hybridization (**Extended Data Fig. 6h**). We also found an expansion of the laminar distribution and the number of BCL11B- and TBR1-immunopositive neurons normally restricted to deep layers, and this may occur at the expense of populations of both SATB2- and CUX1-immunopositive upper layer IT neurons at PD 0 (**Fig. 2j-k, Extended Data Fig. 7d-e**).

To investigate further whether alterations in the expression levels and laminar distribution of specific marker genes associated with IT and ET neurons are correlated with changes in the fate of these neurons, we conducted a series of IdU/CIdU injections at various developmental stages. These injections were administered to both *Neurod6-Cre*; *Zbtb18 cKO* mice and their control littermates. At PCD 12.5, we labeled early-born neurons destined for deep layers (L5-6), while at PCD 14.5 and 15.5, we labeled later-born neurons primarily destined for L4 or L2-3, respectively (**Extended Data Fig. 7a-c**). Subsequently, we analyzed the neocortices of these animals, using immunohistochemistry to access the incorporation of IdU or CldU paired with the expression of specific markers of cell fate, namely, BCL11B for ET neurons and SATB2 for IT neurons. Following the injection of IdU at PCD 12.5, the majority of IdU-positive nuclei were observed in the central region of the CP in both the cKO and control mice, typically where immature L5 neurons are located (**Extended Data Fig. 7a**). While there was a significant increase in the numbers of BCL11B-positive cells and their laminar distribution in the *Neurod6-Cre*; *Zbtb18* cKO mice, there was no corresponding increase in the proportion of cells showing double positivity for IdU and BCL11B immunostaining (**Extended Data Fig. 7a**). This suggests that the laminar identity and position of L5 ET neurons are not substantially altered in mice lacking *Zbtb18* post-mitotically at this age. The later stage injections predominantly labeled upper layer IT neurons that normally express SATB2 but not BCL11B (**Extended Data Fig. 7b-c**). Compared with control animals, in the *Neurod6-Cre; Zbtb18* cKO mice we observed increased proportions of IdU and BCL11B double-labeled nuclei following the injection of IdU at PCD 14.5 (**Extended Data Fig. 7b**). Although the numbers of IdU-positive and BCL11B-positive cells also were greater overall in cKO mice, this suggests that *Zbtb18* post-mitotic deficiency influences cell fate of late-born nascent neurons, aside from any other potential effects on labeled cell populations. Similarly, we observed a decrease in the percentage of nuclei double-positive for CldU and SATB2 following injection at PCD 15.5, explained in part by a broader laminar distribution of the CldU and SATB2 double-positive neurons in the *Neurod6-Cre; Zbtb18* cKO mouse (**Extended Data Fig. 7c**). Collectively, these results indicate that post-mitotic/post-migratory ZBTB18 is required cell-autonomously for proper specification of IT neurons and that its deletion leads to misspecification of at least some IT neurons, including the acquisition of certain molecular properties that are normally associated with deep layer ET neurons.

### ZBTB18 is required for mammalian neocortical projections

The apparent acquisition of ET neuron molecular characteristics by IT neurons in the *Zbtb18* cKO mice suggests that ZBTB18 also may regulate axonal projection development, which is not only a central distinguishing feature of two ExN classes but also a crucial process for the unique mammalian brain connectivity. To investigate the effect of conditional postmitotic *Zbtb18* deletion on the fate of callosal projections, we analyzed early postnatal *Neurod6-Cre Zbtb18* cKO mice that were crossed to mice harboring *CAG-CAT-Gfp*, a CRE-responsive GFP transgene. Consistent with a potential regulatory role in the formation of axonal projection tracts, mutant mice exhibited reduced white matter beneath the subplate and absent CC (**Fig. 3a**). Very few callosal axons reached the midline, and those that did touch the midline then subsequently misrouted ventrally into the septum. Other forebrain commissural projections, including the anterior commissure, were also disrupted, reflecting an overall defect in commissural axon growth (**Fig. 3a**). Additionally, the internal capsule and different sub-cerebral projections also appear to be significantly reduced (**Fig. 3a**).

**Figure 3.**
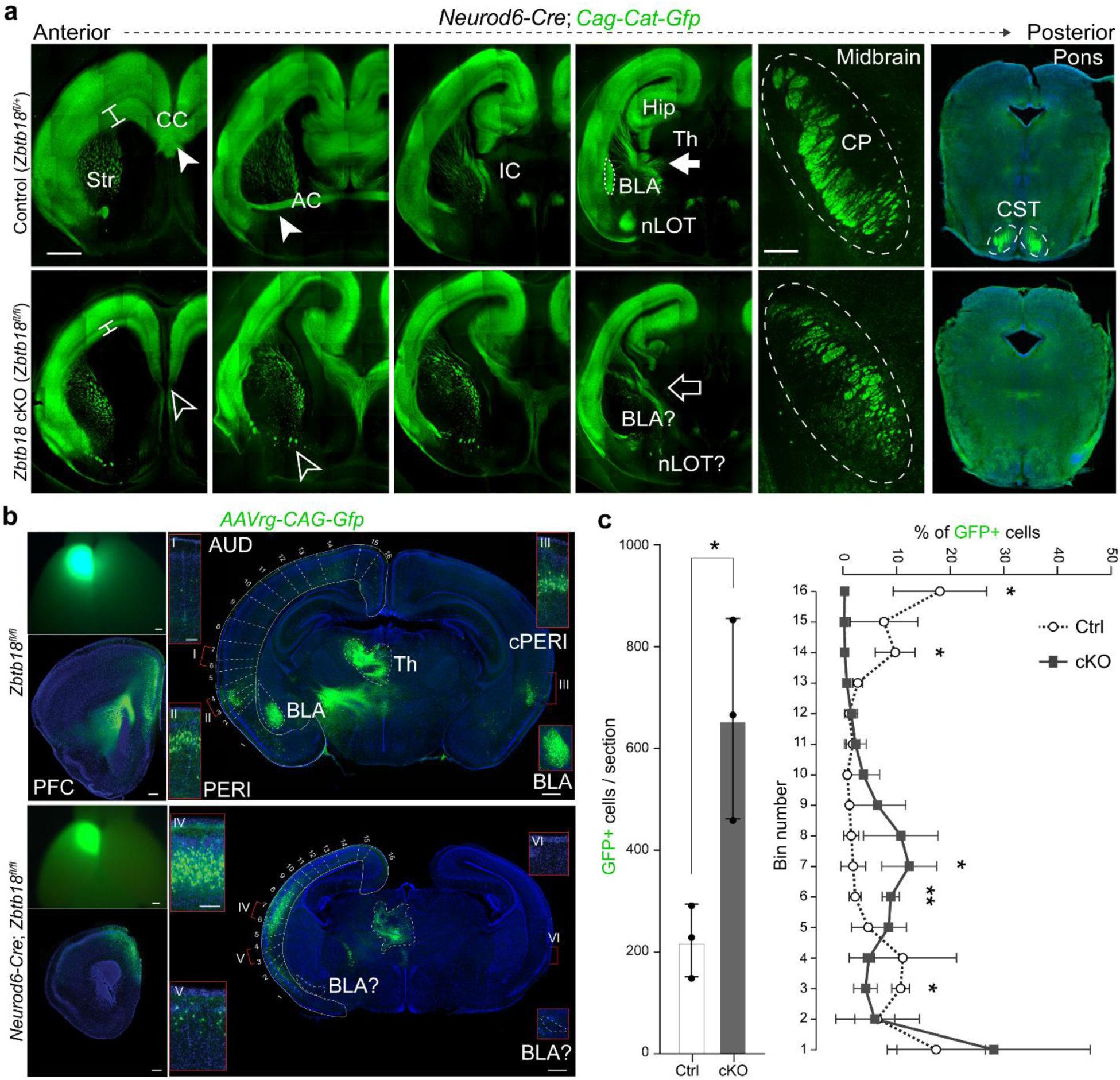
ZBTB18 depletion reduces callosal and subcerebral projections, while increasing cortico-cortical association projections. **a**) Visualization of axonal projections by GFP expression shows, in the *Neurod6-Cre*; *Zbtb18* cKO mouse brain reveals the absence of the CC and CST. Wild-type structures are indicated by full arrows and arrowheads and defective tracts in the *Zbtb18* cKO brain are marked by open arrowheads and arrows. CC, corpus callosum; AC, anterior commissure; Hip, hippocampus; CST, corticospinal tract; Str, striatum; IC, internal capsule; CP, cerebral peduncle; BLA, basolateral amygdala, excitatory nucleus within amygdala; nLOT, nucleus of lateral olfactory tract. Scale bar: 1mm, 20 µm (CP panel). **b**) Whole brain image (left top) and coronal sections of the brain at P7 showing the injection site and extent of the GFP expression after *AAvrg-CAG-Gfp* injections into the medial PFC at PD 3. *AAVrg-CAG-Gfp* was injected into the PFC of PD 3 mice of *Zbtb18* cKO (lower panel) and controls (upper panel) and brains were harvested at PD 7 for the imaging of afferent inputs. Coronal sections of traced PD 7 brains showing more intensive GFP labeling in AUD (bin 6 to 8) than PERI (bin 2 and 3), retrospinal (bin 15 and 16) cortex, and BLA in *Zbtb18* cKO mice than control. No contralateral staining is seen in the cKO mice compared to control mice. Scale bar: 500μm; Inset: 125 μm. PERI, perirhinal cortex; cPERI, contralateral perirhinal; AUD, auditory cortex. **c**) Left: Barplot showing GFP-positive neurons projecting to PFC in the neocortex of Zbtb18 cKO mice compared to control brains at PD 7, as shown in b. unpaired t-test. The graph represents mean ± s.e.m. * P = 0.03 (n = 3). Right: Line graph data showing the percentage distribution of GFP+ neurons projecting to PFC in each bin of neocortex of *Zbtb18* cKO mice compared to wild-type brains at PD 7. The number of labeled neurons in each bin was compared using unpaired t-test. The graph represents mean ± s.e.m. *P = 0.03 (bin3), 0.004031 (bin 6), 0.03 (bin 7), 0.012 (bin 14) (n = 3).

In *Zbtb18* cKOs, we observed a reduction in the long-range inter-cortical and subcortical projections from IT and ET neurons, respectively. To determine whether the intra-hemispheric intracortical connectivity was also affected, we injected *AAVrg-CAG-Gfp*, a GFP-expressing retrograde viral tracing system into the PFC of PD 3 mice. During analysis at PD 7, any brains that did not exhibit proper targeting were excluded to ensure the integrity of the data. In the control group of mice, we observed typical intrahemispheric cortico-cortical connectivity (association pathways) originating from transmodal and unimodal association areas, including the perirhinal cortex, secondary visual cortex, retrosplenial cortex, and parts of the amygdala projecting to the PFC (**Fig. 3b, Extended Data Fig. 8a-b**). However, in the absence of ZBTB18, this pattern of intrahemispheric cortical-cortical association connections appears to be significantly disrupted. Specifically, we noted a substantial increase in unilateral afferent connections to the PFC originating from areas within the central region of the neocortex where, normally, primary areas such as the auditory and somatosensory cortex would be located. Importantly, there was a noticeable reduction in inputs from the perirhinal cortex, secondary visual cortex, and retrosplenial cortex at PD 7 (**Fig. 3b, Extended Data Fig. 8a-b**). Within the neocortex, we also observed an overall increase in the number of GFP-positive cells projecting to PFC (**Fig. 3c**). Additionally, we observed a reduction in the projections from most of the contralateral inputs and subcortical connections as seen in the striatum and thalamus (**Fig. 3b, Extended Data Fig. 8b**). These results suggest compensatory effects after the ZBTB18-mediated loss of callosal and subcortical connections. Collectively, these data demonstrate that ZBTB18 is postmitotically required for the formation of mammalian characteristic neocortical long-range neuronal connectivity.

### ZBTB18 regulates mammalian *Robo1* enhancer and callosal formation

To examine whether ZBTB18 regulates callosal projections in a cell-autonomous manner, next we co-electroporated the *Neurod1-Cre* and *CALNL-Gfp* reporter vectors into the neocortex of PCD 15 *Zbtb18^fl/fl^* mice. This *Cre*-mediated deletion of *Zbtb18* from ExNs resulted in their diminished ability to produce GFP-labeled axons projecting towards or across the midline at PD 0 (**Fig. 4a**). Co-electroporation with a *Neurod1-Zbtb18* expression vector that restores ZBTB18 in the same cell populations, however, rescued the projection of GFP-positive callosal axons to the midline (**Fig. 4a**). These results demonstrate that ZBTB18 is cell-autonomously required for the midline crossing of callosal axons.

**Figure 4.**
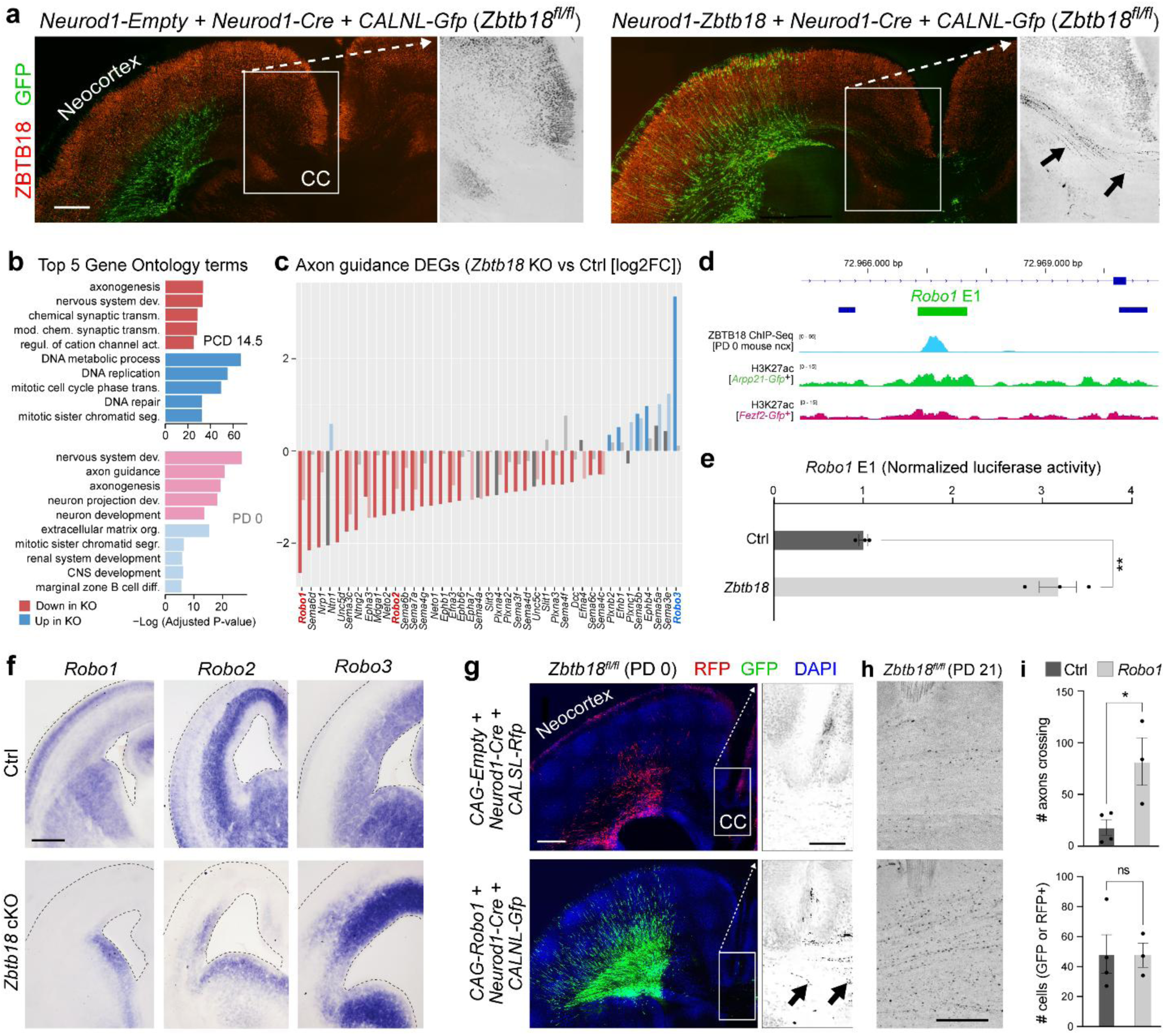
ZBTB18 regulates axon guidance genes and the mammalian-specific *Robo1* enhancer. **a**) Left panel: Representative images of *in utero* electroporation of *Neurod1-Cre* and *CALNL-Gfp* plasmids into *Zbtb18* ^fl/fl^ mice with double immunofluorescence shows *Zbtb18* is required for the formation of the CC. Right panel: Representative images of co-electroporation of the *Neurod1-Zbtb18* constructs rescued this callosal projection phenotype. Scale bar: 500 μm. **b**) Top 5 Gene ontology terms for the genes that are downregulated (red) or upregulated (blue) in the *Zbtb18* -/- (KO) mouse as compared to *Zbtb18 +/-* (control) at PCD 14.5 (upper bar plot) and in *Neurod6-Cre*; *Zbtb18* cKO as compared to control at PD 0 (lower bar plot). **c**) Genes encoding axon guidance molecules and up-and down-regulated in the *Zbtb18* KO mouse as compared to *Zbtb18 +/-* (control). **d**) Line graphs showing the H3K27ac peaks from *Arpp2-Gfp*+ IT neurons, *Fezf2-Gfp*-positive ET neurons and ZBTB18-HA ChIP-Seq peaks associated with mouse *Robo1*. **e**) Luciferase reporter activity driven by the *Robo1* E1 enhancer is significantly increased by ZBTB18. An unpaired t-test was used to detect differences between control and experimental conditions. The graph represents mean ± s.e.m. ** P = 0.00672 (n = 3). **f**) *In situ* hybridization shows the reduction of *Robo1* and *Robo2* expression in the neocortical plate and the induction of *Robo3* expression in the *Zbtb18* -/- mouse. Scale bar: 150 μm. **g-i**) Representative images of the *in utero* electroporation of a *Robo1* expression plasmid (*CAG-Robo1*) allow Gfp-expressing, co-electroporated upper layer neurons to project GFP-positive axons to and across the CC (arrows in the inset) analyzed at PD 0 (g) and PD21 (h); Upper graph depicts the number of axons crossing at the midline on the contralateral side of the IUE at PD 21 (i) and the lower graph shows the number of cells electroporated on the ipsilateral side (i). An unpaired t-test was used to detect differences between control and experimental conditions. The graph represents mean ± s.e.m. * P = 0.030. Scale bar: 500μm, inset 250μm. For each *in utero* electroporation experiment, 6 control and 3 *Robo1* successfully electroporated animals were analyzed at PD 0; at PD 21, 3 control and 3 *Robo1* successfully electroporated animals were analyzed. RNA-Seq was conducted on at least 3 independent biological replicates for each condition. *In situ* hybridization data shown are representative of data generated from multiple sections of at least 3 animals.

We next sought to understand the potential downstream effectors of ZBTB18 in this process by analyzing our neocortical RNA-seq data from *Zbtb18* KO mice and littermate controls. As expected, gene ontological terms associated with up- and down-regulated genes at both PCD 14.5 and PD 0 referenced general categories such as nervous system development, and several terms implicating DNA replication and the cell cycle were enriched among upregulated genes at PCD 14.5. Ontology terms, including axonogenesis, axon guidance, and neuron projection development, were also enriched among genes downregulated in the ZBTB18 knockout at both PCD 14.5 and PD 0, consistent with a role for ZBTB18 in regulating neuronal projection formation (**Fig. 4b**). These observations suggest that ZBTB18 not only promotes cell cycle exit and neuronal migration, as previously described ^13,16,17^, but may also promote postmitotic neocortical ExN diversification and axonogenesis through the regulation of genes involved in IT neuron fate commitment and projection system development.

To explore the molecular underpinnings for the failure of appropriate callosal axon targeting by *Zbtb18*-deficient projection neurons, we analyzed the RNA-Seq data for changes in the expression of genes encoding axon guidance and cell adhesion molecules. We found that the expression of several receptors previously implicated in callosal development, such as members of the Netrin (i.e., *Dcc* and *Unc5d* ^35–37^) and Slit-Robo families of axon guidance molecules were either downregulated (*Robo 1* and *2* ^38,39^) or upregulated (*Robo 3* ^40^) in the *Zbtb18* KO neocortex (**Fig. 4c**). Notably, the Slit receptor, *Robo1*, was the most downregulated in the *Zbtb18* KO neocortex. Additionally, ChIP-Seq data performed against HA-tagged ZBTB18 revealed an IT neuron-specific putative enhancer, *Robo1* E1, within the *Robo1* locus (**Fig. 4d, Extended Data Fig. 9a**). Luciferase activity assays demonstrated significant activation of *Robo1* E1 by ZBTB18 as compared to controls (**Fig. 4e**). *In situ* hybridization at PCD 15.5, the period when callosal projections are typically being formed in mice, confirmed the downregulation of *Robo1* and *Robo2*, as well as the upregulation of *Robo3* in the *Zbtb18* cKO neocortex (**Fig. 4f**).

Given that *Robo1* emerged as the significantly downregulated axon guidance gene within the *Zbtb18* KO neocortex, attributable to its direct regulation by ZBTB18, and considering the observed callosal defects resulting from disruptions in both mouse and human homologs ^38,39,41^, sought to ascertain any potential correlation between alterations in Robo1 expression and the manifested callosal projection anomalies. For this, we electroporated a control (*CAG-empty*) or *Robo1*-expression vector (*CAG-Robo1*) together with *Neurod1-Cre* and *CALNL-Gfp* (or *CALNL-Rfp*) into the PCD 15.5 *Zbtb18^fl/fl^* mouse neocortex. As expected, we found negligible numbers of labeled axons projecting towards or across the midline at PD 0 in any of six control electroporated *Zbtb18^fl/fl^* brains (**Fig. 4g**). In contrast, we observed the recovery of a proportion of GFP+ axons in the CC following *Robo1* over-expression in all electroporated animals (**Fig. 4g**). To test the robustness and persistence of such changes, we then examined the rescue of the CC at a later stage when its development is complete. Consequently, we conducted analyses at PD 21 and found similar phenotypic rescue as in PD 0 brains. Specifically, we observed the restoration of a subset of GFP+ axons in the CC upon overexpression of *Robo1* (**Fig. 4g-i**). The fact that the observed *Robo1*-mediated recovery of callosal projections was not extensive suggests that midline crossing is a complex cellular and molecular process and that other factors, such as additional cell-intrinsic effectors (e.g., DCC and ROBO2), or the actions of midline glia ^8^, may be required for callosal projections or a combination of these (especially in humans and in relation to pathology).

### High conservation of ZBTB18 binding sites in mammalian IT CREs

The finding that ExN subtype-specific CREs controlled by ZBTB18 display either mammal-specific characteristics or possess unique mammalian ZBTB18 binding sites led us to explore whether the evolution of this regulatory system might mark a significant evolutionary milestone in mammals. To assess this possibility, we first analyzed the protein coding sequences and dorsal pallial expression of *Zbtb18* and found that they are highly conserved across vertebrates (**Fig. 1h-i, Extended Data Fig. 1e, Extended Data Fig. 10a**). Next, we considered the conservation of broad populations of ZBTB18 binding motifs across placental mammals, marsupials, monotremes, and non-mammalian species. Then we assessed the evolutionary footprint of specific CREs and/or ZBTB18 binding motifs plausibly regulating key genes associated with major subtypes of ExNs, including *Cux2*, *Satb2, Bcl11b*, *and Robo1*. However, because variations in TF binding sites may lead to interspecies differences ^23,24,26,27^, we sought to determine whether variations among putative ZBTB18 binding sites are conserved within specific clades. We therefore assessed the relative conservation of putative ZBTB18 binding sites among CREs we identified near genes whose expression was enriched in IT neurons (**Extended Data Fig. 10a**), and we compared this to CREs unrelated to IT/ET neuron subspecification or CC formation (see methods). In addition, we considered CREs near genes whose expression was enriched in ET neurons or whose expression level was up- or down-regulated in *Zbtb18* KO mice. When precise matching with the core motif as described in the JASPAR database (**Extended Data Methods**) was required, we found ZBTB18 binding sites were significantly conserved in IT neuron enhancers in placental mammals and marsupials but not in monotreme or non-mammalian species (**Fig. 5b**). This enrichment was not observed in ET neuron enhancers. Moreover, we found significant conservation of ZBTB18 binding sites in the CREs of genes we observed to be upregulated in *Zbtb18* KO mice specifically throughout Eutheria but no other mammals (**Fig. 5a**). We also assessed the level of conservation of ZBTB18 binding sites with 1, 2, or 3 mutations in the core sequence, reasoning that heavily mutated, and thus likely non-functional, ZBTB18 binding sites should not exhibit similar enrichment in conservation. Expectedly, we found no enrichment among putative IT neuron-associated enhancers or those upregulated in *Zbtb18* KO mice when we assessed conservation with either 2 or 3 mutations, which suggests the ZBTB18 binding site conservation is specific among IT neurons (**Fig. 5a**).

**Figure 5.**
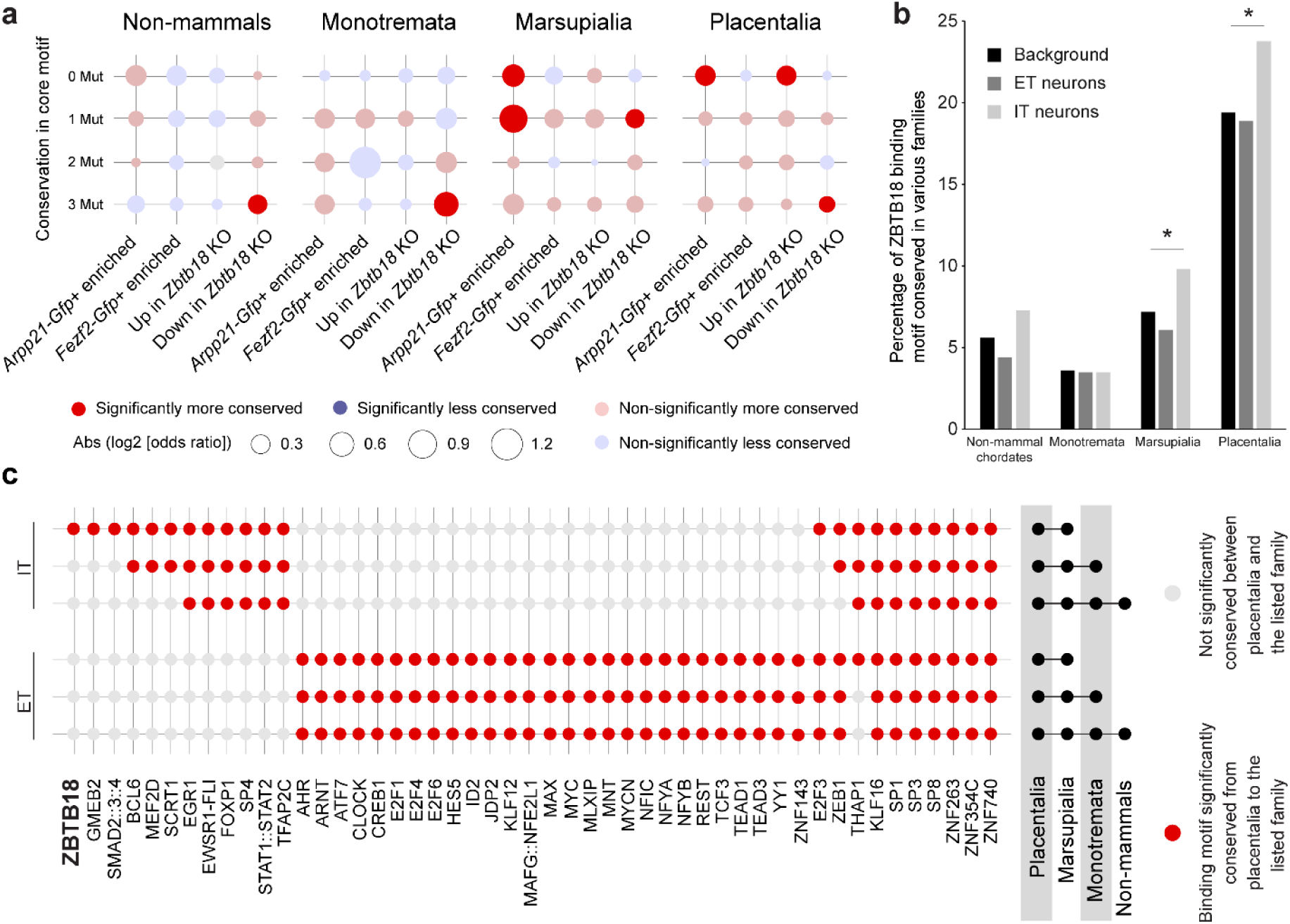
Increased conservation of ZBTB18 binding motifs in mammalian IT neuron-biased CREs. **a**)Conserved ZBTB18 binding motifs are identified in putative CREs associated with genes exhibiting enriched expression in *Arpp21-Gfp*+ IT neurons. **b**) Between marsupials and placental mammals, enhancers associated with *Arpp21-Gfp*+/IT neurons have a significantly higher percentage of conserved ZBTB18 motifs than those associated with *Fezf2-Gfp*+/ET neurons or background sequences. No differences of ZBTB18 motif conservation in non-mammals or monotremes were detected. Asterisks indicate where Fisher’s exact test revealed a significant enrichment of conservation, with a False Discovery Rate corrected P-value < 0.05. **c**) Among the TFBS cataloged in the JASPAR database and expressed in either IT neurons or ET neurons, the ZBTB18 motif stands out as one of three motifs with consensus sequences uniquely conserved within enhancers linked to genes enriched in IT neurons across placental mammals and marsupials. Fisher’s exact test was used to test for an enrichment of conservation. Red dots represent motifs with a False Discovery Rate corrected P-value < 0.05.

Considering the possibility that this high degree of conservation in placental mammals and marsupials may not be specific just for ZBTB18, we next performed a similar analysis using 267 TF binding motifs from the JASPAR database, corresponding to TFs that are expressed in either IT- or ET-neurons, or both, at PD 0 according to our RNA-Seq dataset. We again assessed for the enrichment of binding sites with perfect core motif conservation in CREs associated with IT- or ET-neuron enriched genes or those differentially regulated in the *Zbtb18* KO (**Fig. 5c**). We grouped the significant motifs according to independently tested enrichment in consecutive phylogenetic groups (i.e., Placentalia + Marsupalia, Placentalia + Marsupalia + Monotrema, etc). Of the 267 motifs, only two other motifs (i.e., glucocorticoid modulatory element binding protein 2 (GMEB2) and SMAD2/3/4, involved in transforming growth factor (TGF)-beta signaling) in addition to ZBTB18 were found to be specifically conserved in IT neuron-associated CREs in Theria, demonstrating that ZBTB18 is a member of a rare group of TFs, including the two involved in glucocorticoid or TGF-beta signaling, with binding sites that are singularly conserved among placental mammals and marsupials in IT neuron-related CREs (**Fig. 5c**). This further supports a critical role for ZBTB18 in the evolution of the mammalian diversification of ExNs. Interestingly, among genes significantly upregulated in the neocortex of *Zbtb18* KO mice, the enriched conservation of ZBTB18 binding motifs was only significant across placental mammals (Eutheria) (**Extended Data Fig. 10b**).

Finally, to probe the evolutionary origin of *ZBTB18* genes, we estimated their evolutionary age by protein sequence similarity and found that, while *ZBTB18* genes are ancient and shared with bacteria, their protein sequences do not change amongst chordates (**Extended Data Fig. 10a**). Collectively, these results suggest that it may not be the expression or the protein structure and organization of ZBTB18 itself but the presence of the evolutionary-derived CREs that is essential to drive the expression pattern of key genes involved in the molecular sub-specification and connectivity of neocortical ExNs within mammals.

## Discussion

This research addressed a fundamental question at the intersection of neuroscience and evolutionary biology: the emergence of the mammalian neocortex, particularly its diverse ExN subtypes and their intricate long-range projections. Our findings revealed and characterized mammalian-specific adaptations at the level of CREs linked to TFs most associated with ET/IT neuron specification. These adaptations contribute to the evolutionary divergence observed in mammalian CREs, gene expression patterns, ExN subtype specification, and long-range projections compared to non-mammalian counterparts. We also revealed a key subgroup of CREs associated with IT and ET neuron specifying genes either exhibit mammalian-specific characteristics or contain unique ZBTB18 binding sites exclusive to mammals. We also show that certain mammalian-specific changes associated with ZBTB18 within those CREs lead to uniquely mammalian repression or activation of genes essential for proper molecular diversity and connectivity of mouse ExNs. We found that ZBTB18 is a highly conserved TF with regard to its structure as well as its high expression level in the cortex. However, CREs evolve at a much faster speed, and rapid enhancer evolution is a fundamental trait of mammalian genomes ^24,26^. Therefore, this evolutionary-developmental aspect of our study revealed the critical elements and evolutionary adaptations within the gene regulatory network that underpin the unique features of mammalian neocortical ExNs.

Consistent with these findings, the loss of ZBTB18 function in mice leads to simplified neocortical laminar organization and long-range axonal projection patterns that resemble ancestral forms of laminar and intrahemispheric cortico-cortical connectivity. This research also expands upon the study of ExNs in the mouse piriform cortex (paleocortex) that revealed diminished ExN subspecification compared to the six-layered neocortex, with piriform ExNs retaining molecular signatures reminiscent of ancestral cortical cell identities shared with reptiles and amphibians ^42^. Interestingly, the expression level of *Zbtb18* in immature ExNs of the piriform cortex is substantially lower than in the neocortex (**Extended Data Fig. 4a**), suggesting a potential link between *Zbtb18* expression and ExN diversity across the paleocortex and neocortex.

Human genetic studies have associated the loss of function ZBTB18 variants with agenesis of the CC, microcephaly, autism, intellectual disability, and related phenotypes ^20–22,43,44^. In addition, mutations in here identified target genes of ZBTB18 that have undergone mammalian-specific changes in their CREs and expression patterns, which include *CUX2*, *SATB2*, and *BCL11B*, have also been linked to intellectual disability and autism ^21,45–47^. Our research reveals that the CREs and cell-type specific expression patterns of these disease-associated genes within the neocortex are highly conserved across mammalian species and differ markedly from that seen in non-mammalian counterparts. Furthermore, ZBTB18 binding sites are highly conserved in Eutherian IT-biased CREs.

Consequently, this heightened evolutionary constraint on this crucial regulatory node in the mammalian neocortex, while advantageous, may also render them more susceptible to various neurodevelopmental and neuropsychiatric disorders, where deviations from the typical circuits are often noted. Furthermore, we provide potential mechanistic links between dysfunction in this important regulatory node and the potential future utility in the functional interpretation of human genetic variants within the characterized CREs related to neocortical phenotypes observed in affected individuals. However, we don’t know if variations in any of the identified CREs are associated with intellectual disability or autism. Furthermore, our research has demonstrated potential predictive value for understanding the pathogenesis of autism. Our findings indicate that individuals with ZBTB18 loss of function mutations or 1q43q44 microdeletion syndrome, where ZBTB18 haploinsufficiency plays a significant role, might exhibit increased interhemispheric cortico-cortical connectivity between the PFC and temporal heteromodal association areas. This might also extend to other cases of intellectual disability and autism. Supporting this hypothesis, resting-state functional magnetic resonance imaging (fMRI) studies have observed increased functional connectivity, primarily involving PFC and temporal heteromodal association areas in the default mode network and hypoconnectivity in sensory-motor and visual areas in autism ^48–50^. These dysconnectivity patterns are strongly correlated with behavioral symptoms observed in autism ^50^. However, we don’t know if the increased interhemispheric mPFC-temporal association projections in *Zbtb18* cKO are also associated with enhanced functional connectivity. While we did not directly examine connectivity within sensory-motor and visual networks, due to an apparent expansion of temporal heteromodal projections in the *Zbtb18* cKO mice, we predict that we would observe smaller and interhemispherically hypoconnected sensory-motor and visual areas in these mice. In conclusion, our findings underscore the relevance of these evolutionary and developmental insights in the translational context, as alterations in this specific regulatory node are linked to major neurodevelopmental and neuropsychiatric disorders.

## Acknowledgments

We thank members of the Heng, Richards, and Sestan labs for their extremely helpful comments. We also wish to thank Constance Cepko, Elizabeth Grove, Franck Polleux, and Pasko Rakic for providing reagents. This work was supported by the Australian Research Council (DP 160103958 to R.S. and L.J.R., and Discovery Early Career Researcher Award Fellowship 160101394 to R.S.), the National Health and Medical Research Council (Principal Research Fellowship to L.J.R., and C.J. Martin Fellowship to P.K.), the Australia Postgraduate Award (L.R.F.) and the UQ-QBI Doctoral Scholarship (A.P.), the Victorian Government through the Operational Infrastructure Scheme and the National Health and Medical Research Council of Australia (ID:1011505 to J.I.H.), and Instituto de Salud Carlos III Spain and European Social Fund grant MS20/00064 (to G.S.); grant PID2019-104700GA-I00 and PID2022-140137NB-100 funded by /AEI/10.13039/501100011033 (to G.S.); and Fundació LaMarató de TV3(to G.S); and NIH grants MH129981 (to K.T.G) and NS095654, MH106934, MH116488, and MH110926, MH122678, MH124619, HG010898 and the Simons Foundation 736706 (to N.S.).

## Authors Contributions

Z.L., N.K., S.K.S., C.Q. and M.S. performed a majority of the genetic, molecular, and other experiments described in this study. S.K.M., G.S., X.D.M. performed a majority of the computational, bioinformatic, and related analyses for this study. Additionally, M.L., X.X., P.K., R.S., L.V., A.K., performed additional computational analysis. Z.L., O.C., A.S., I.K.S., T.H., and F.O.G. conducted *in situ* hybridization experiments. Z.L., N.K., S.S., A.M.M.S., R.D.R., D.P., A.T.N.T., T.K., W.H., A.P., and L.R.F. performed analysis of immunohistochemistry or protein content. Z.L., W.H., S.K.S, and Y.B. performed *in utero* electroporation experiments. Z.L., A.T.N.T., and S.P. developed/acquired human tissue resources used in this study. Z.L., X.X., E.G-N., and H.D.C. developed/acquired murine model resources used in this study. Z.L., S.K.M., G.S., F.O.G., and N.S. prepared and wrote the initial draft of this manuscript, and all authors provided edits and/or comments on subsequent drafts. Q.Z., F.O.G., N.K., A.M.M.H, L.R., J.I.H., and N.S. provided supervision and guidance for this study. N.S. conceived and designed the study. L.R., J.I.H., G.S. and N.S. obtained and provided funding for these efforts.

