## Supplementary Figures for "Adaptive Evolution of Gene Regulatory Networks in Mammalian Neocortical Neurons"

### Extended Data Figures

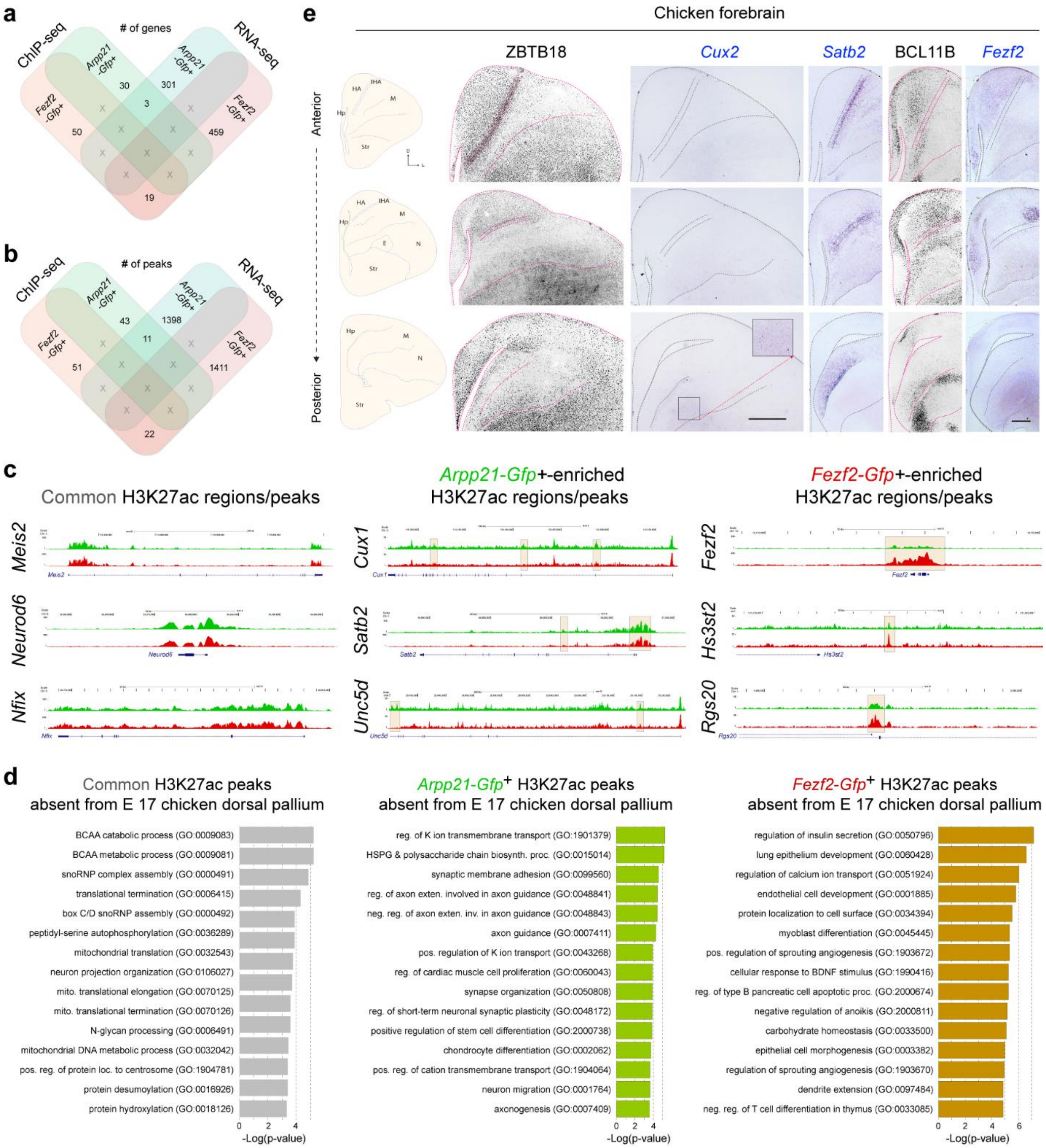


**Extended Data Figure 1. Integration of RNA-Seq and ChIP-Seq to identify gene enrichment and CREs in IT and ET neurons**

**a-b**) Venn diagrams illustrating the overlap between genes differentially expressed and genes regulated by differentially enriched H3K27ac peaks between *Arpp21*- and *Fezf2*- labeled cells. **b**)

As in A but representing overlap of H3K27ac peaks rather than genes. **c**) Most genes and H3K27ac peaks were not differentially expressed or enriched, respectively, between IT neurons and ET neurons (left panel). However, certain genes were enriched in either IT neurons or ET neurons and associated with putative CREs enriched or present in that same cell subtype (middle and right panels). Data shown represent ChIP-Seq data collected from 2 independent biological replicates per condition. **d**) Bar graph showing the top 15 gene ontology terms in the H3K27ac peaks either present in both (grey) or in *Arpp21-Gfp*+ (green) or in *Fezf2-Gfp*+ (red) cells but are absent in the H3K27ac peaks from E 17 chicken. The X-axis depicts the log2 P-value. **e**) Schematics showing different areas of the chicken brain across the anterior-posterior axis in the coronal plane. Coronal sections of chicken brain showing ZBTB18, *Cux2*, *Satb2*, BCL11B, and *Fezf2* expression domain ZBTB18 and BCL11B images showing ZBTB18 and BCL11b immunostaining are inverted to enhance visualization. In inset *Cux2* in situ expression signal is shown at posterior plane. This demonstrates the *Cux2* probe's sensitivity to chicken *Cux2*. Scale bar: ZBTB18, *Cux2*, (500 µm); *Satb2*, BCL11B*, Fezf2* (250 µm). Hp, hippocampus; HA, apical hyperpallium; IHA, interstitial apical hyperpallium, M, mesopallium; N, nidopallium; E, entopallium; Str, striatum; D, dorsal; L, lateral.

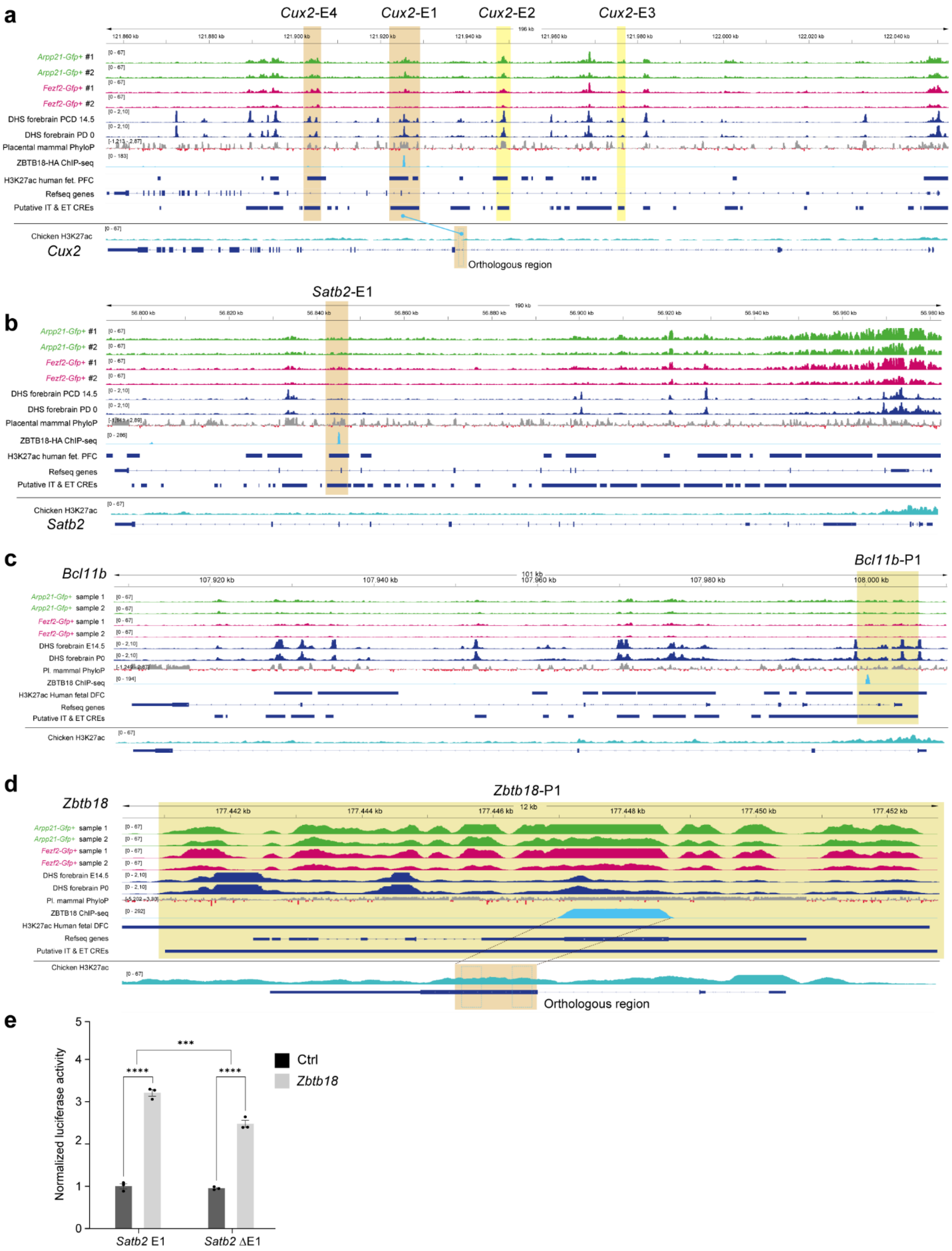
a
Cux2-E4
Cux2-E1
Cux2-E2
Cux2-E3
Arpp21-Gfp+ #1
Arpp21-Gfp+ #2
Fezf2-Gfp+ #1
Fezf2-Gfp+ #2
DHS forebrain PCD 14.5
DHS forebrain PD 0
Placental mammal PhyloP
ZBTB18-HA ChIP-seq
H3K27ac human fet. PFC
Refseq genes
Putative IT & ET CREs
Chicken H3K27ac
Cux2
Orthologous region
b
Satb2-E1
Arpp21-Gfp+ #1
Arpp21-Gfp+ #2
Fezf2-Gfp+ #1
Fezf2-Gfp+ #2
DHS forebrain PCD 14.5
DHS forebrain PD 0
Placental mammal PhyloP
ZBTB18-HA ChIP-seq
H3K27ac human fet. PFC
Refseq genes
Putative IT & ET CREs
Chicken H3K27ac
Satb2
c
Bcl11b
Bcl11b-P1
Arpp21-Gfp+ sample 1
Arpp21-Gfp+ sample 2
Fezf2-Gfp+ sample 1
Fezf2-Gfp+ sample 2
DHS forebrain E14.5
DHS forebrain P0
Pl. mammal PhyloP
ZBTB18 ChIP-seq
H3K27ac Human fetal DFC
Refseq genes
Putative IT & ET CREs
Chicken H3K27ac
d
Zbtb18
Zbtb18-P1
Arpp21-Gfp+ sample 1
Arpp21-Gfp+ sample 2
Fezf2-Gfp+ sample 1
Fezf2-Gfp+ sample 2
DHS forebrain E14.5
DHS forebrain P0
Pl. mammal PhyloP
ZBTB18 ChIP-seq
H3K27ac Human fetal DFC
Refseq genes
Putative IT & ET CREs
Chicken H3K27ac
Orthologous region
e
Normalized luciferase activity
Ctrl
Zbtb18
****
****
***
Satb2 E1
Satb2 ΔE1

**Extended Data Figure 2**. **CREs bound by ZBTB18 at the genomic loci of *Cux2, Satb2, Bcl11b* and *Zbtb18* and luciferase reporter assay**

**a-d**) Line graphs showing the H3K27ac peaks from, *Arpp21-Gfp*+ cells (line 1 and 2), *Fezf2-Gfp*+ cells (line 3 and 4), from PCD 14.5, and PD 0 forebrain DNaseI Hypersensitivity tracks (DHS) by Digital DNaseI from ENCODE, (line 5 and 6, respectively) and mammal conservation as reference (line 7), ZBTB18-HA ChIP-seq peaks from mouse (line 8), H3K27ac peaks from fetal (dlPFC (line 9), gene locus in mice (line 10) H3K27ac peaks collapsed together in the *Arpp21-Gfp*+ IT neurons and *Fezf2-Gfp*+ ET neurons (line 11) and H3K27ac peaks from E17 chicken dorsal pallium (12) at the genomic locus of *Cux2, Satb2*, *Bcl11b* and *Zbtb18*. The H3K27ac and ZBTB18- ChIP peaks in mice are shown in light beige. In the chicken genome, the regions orthologous to the ZBTB18-ChIP peaks are shown in a dark beige color, with the dotted blue boxes indicating the exact orthologous regions obtained after liftover. **e**) Luciferase reporter activity driven by the *Satb2* E1 enhancer is significantly increased by ZBTB18 and reduces when the ZBTB18 binding site is mutated (*Satb2* ΔE1). Ordinary two-way ANOVA with Bonferroni's multiple comparisons test, with single pooled variance, was applied. The graph represents mean ± s.e.m. **** P = 0.0001 (*Satb2* E1 (Control vs *Zbtb18*)), (*Satb2* ΔE1 (Control vs *Zbtb18*)), and P = 0.003 for *Satb2* E1 vs *Satb2* ΔE1 (*Zbtb18*), n = 3.

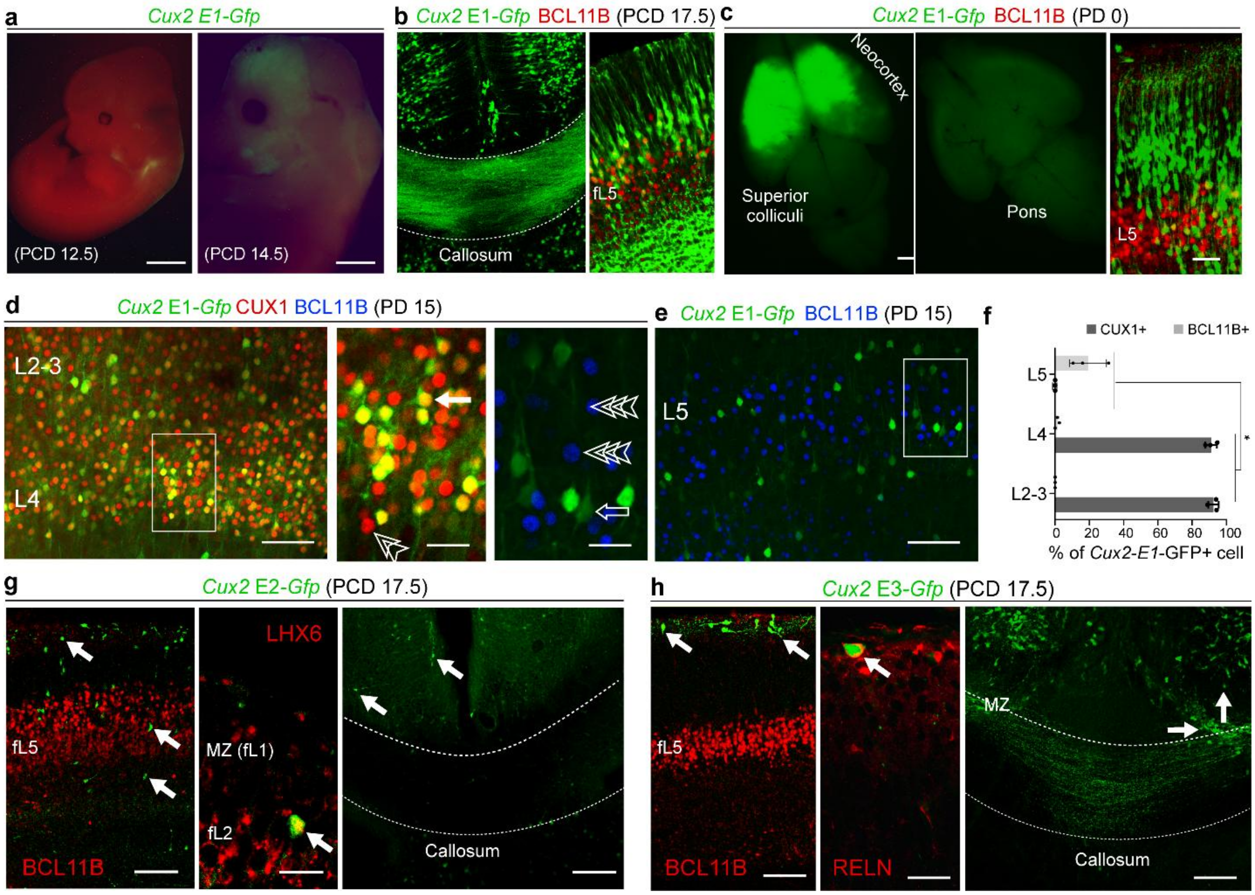


**Extended Data Figure 3**. **Assessment of Cux2 enhancer activities through mouse transgenic reporter assays**

**a**) Transgenic mouse *Cux2-E1-Gfp* shows no GFP expression at PCD 12.5, but strong GFP expression at PCD 14.5 GFP in the embryonic mouse forebrain. Scale bar: 1mm (left), 2mm (right). **b**) At PCD 17.5, GFP is expressed in the BCL11B negative cells in upper CP and CC in *Cux2-E1-Gfp* mice. Scale bar: 1mm (whole brain), 50 µm (right). **c**) At PD 0, GFP is mostly expressed in BCL11B negative upper layer neurons and absent from CST as seen at pons in *Cux2-E1-Gfp* mice. Scale bar: 1mm (whole brain), 50 µm (right). (**d-f**) At PD 15, the majority of *Cux2-E1-Gfp* positive cells reside in upper layers and co-label with CUX1 (closed arrows). Double open arrowheads (CUX1) and triple open arrowheads (BCL11B) again indicate cells immunolabeled for specific molecular markers but not by *Cux2-E1-Gfp.* CUX1 and BCL11B positive cell populations were compared using an unpaired t-test. The graph represents mean ± s.e.m. * P = 0.000352. Scale bars: 100µm (d); inset, 50µm;100µm (e). **g**) In the PCD 17.5, *Cux2-E3-Gfp* transgenic mice brain, GFP is mostly expressed in L1 cell and co-localize with RELN, but not with BCL11B, GFP labeled axons can be seen passing through CC. Scale bar: 100 µm (left, right), 20 µm (middle). (**h**) In the PCD 17.5 *Cux2-E2-Gfp* transgenic mice brain, GFP-positive cells are scattered throughout CP and largely co-localize with LHX6; GFP is not expressed in CC. Scale bar: 100 µm (left, right), 20 µm (middle).

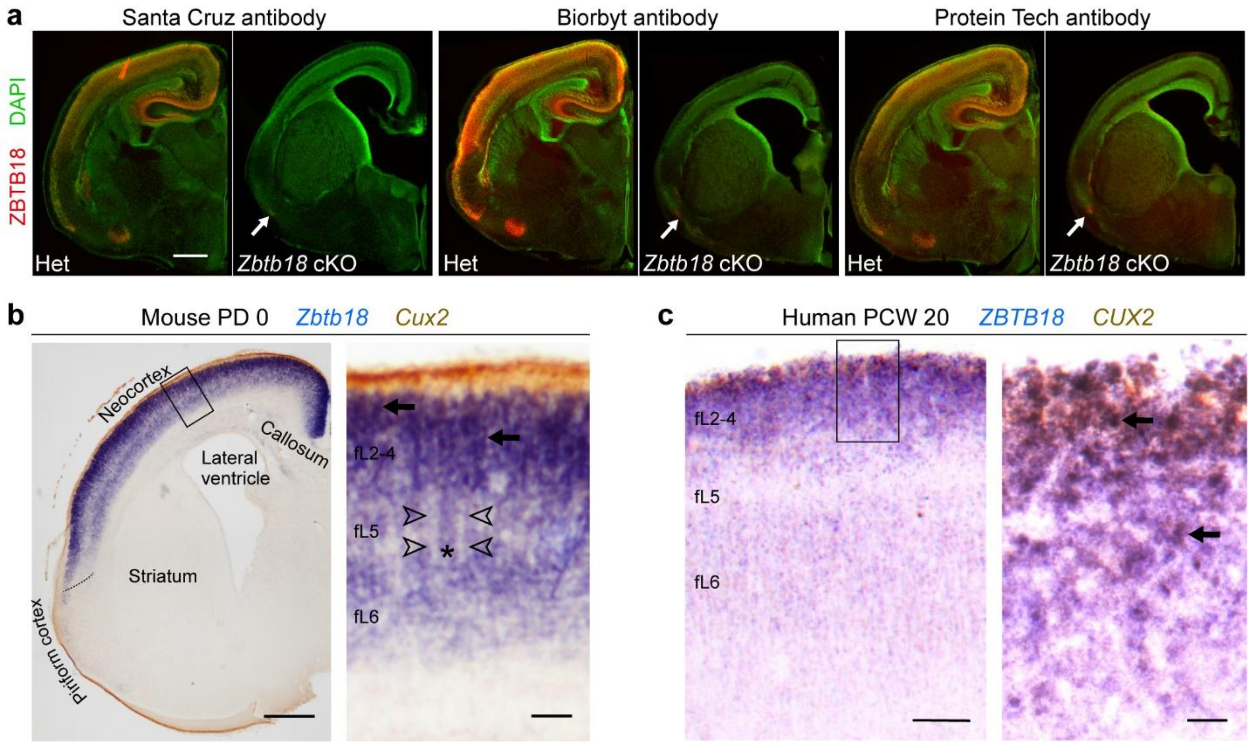


**Extended Data Figure 4**. **ZBTB18 expression in the developing cerebral cortex of humans and mice**

**a**) Coronal sections showing the immunolabeling for three anti- ZBTB18 antibodies on *Neurod6-Cre; Zbtb18* cKO as compared to the control. Arrow points to the expression of ZBTB18 in the ventral region of the cKO where *Neurod6-Cre* appears not be active. Scale bar: 150 µm. **b**) Representative images of double-labeled in situ hybridization in a postmortem PD 0 mouse forebrain reveal striking co-expression of *Zbtb18* and *Cux2* mRNAs in fetal/neonatal layers 2 to 4 (fL2-4), as indicated by arrows. Notably, within the fL5, *Zbtb18* exhibits robust expression in select cells, denoted by asterisks, while other cells in this layer show comparatively lower expression levels, marked by open arrowheads. Scale bar: 1 mm (left), 20 µm (right). **c**) Representative images of double-labeled in situ hybridization in a postmortem postconception weeks (PCW) 20 human frontal cortex reveal co-expression of *ZBTB18* and *CUX2* mRNAs (arrow). Scale bar : 2 mm, 40 µm (right) CP, cortical-plate ; IZ, intermediate zone; SVZ, subventricular zone; VZ, ventricular zone. All images of mouse neocortex are representative of at least 3 independent biological replicates. *In situ* hybridization of PCW 20 human neocortex reflects multiple sections from a single brain but is consistent with an additional PCW 18 postmortem human brain sample (not shown).

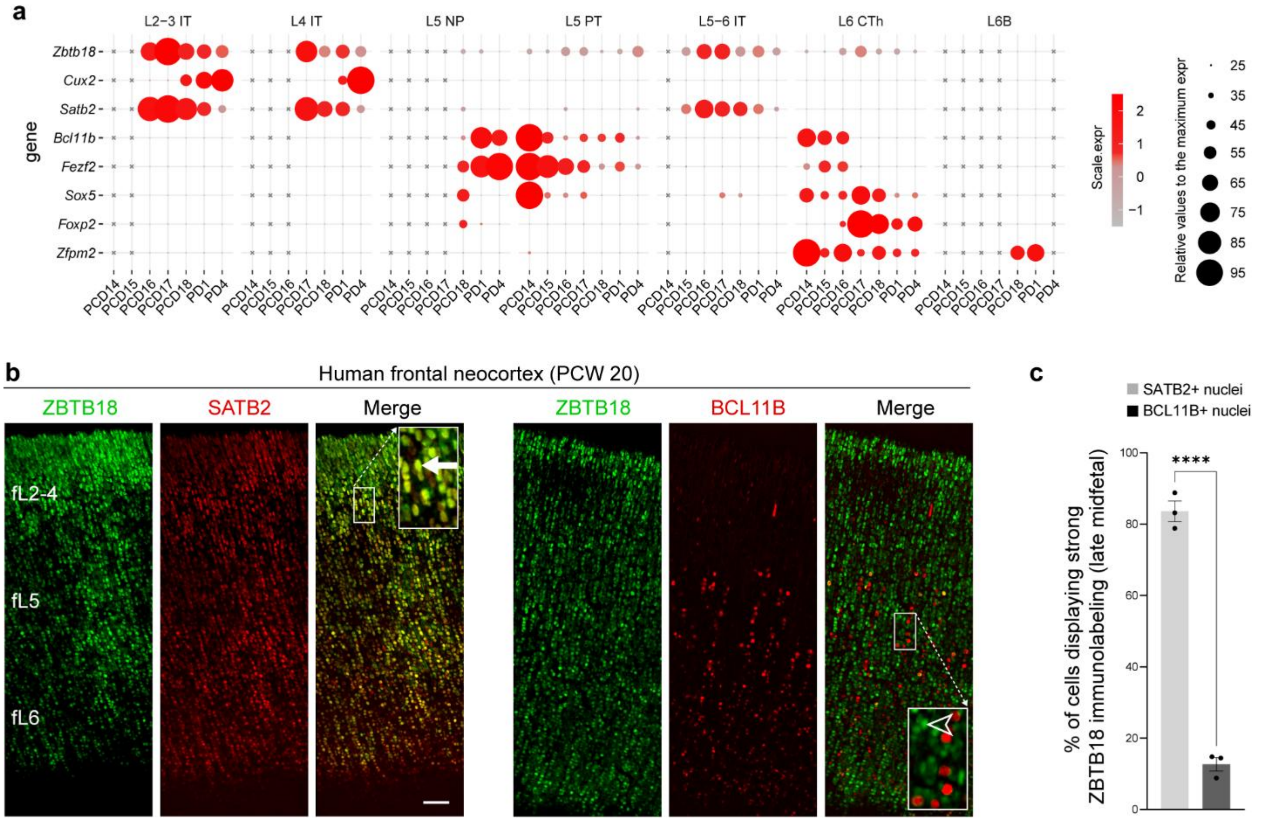


### Extended Data Figure 5. Enrichment of ZBTB18 in postmigratory IT neurons

**a**) Dot-plot showing expression of *Zbtb18* and other canonical layer markers across ExN types along developmental ages of mouse cortex [34]. *Zbtb18* co-expresses *Satb2* and *Cux2* in IT neurons but not with ET neurons' TFs *Bcl11b*, *Fezf2*, *Sox5*, *Foxp2*, and *Zfpm2*. **b-c**) ZBTB18 co-localizes with SATB2, but not BCL11B, in the 20 PCW human neocortex. Scale bar: 100µm. Immunofluorescent analysis of the mouse brain utilized independent sections from at least 3 animals. Images and quantification shown in A are representative of analysis conducted on at least 2 sections from the postmortem human brain. An unpaired t-test was applied. The graph represents mean ± s.e.m. * P = 0.0001, n = 3.

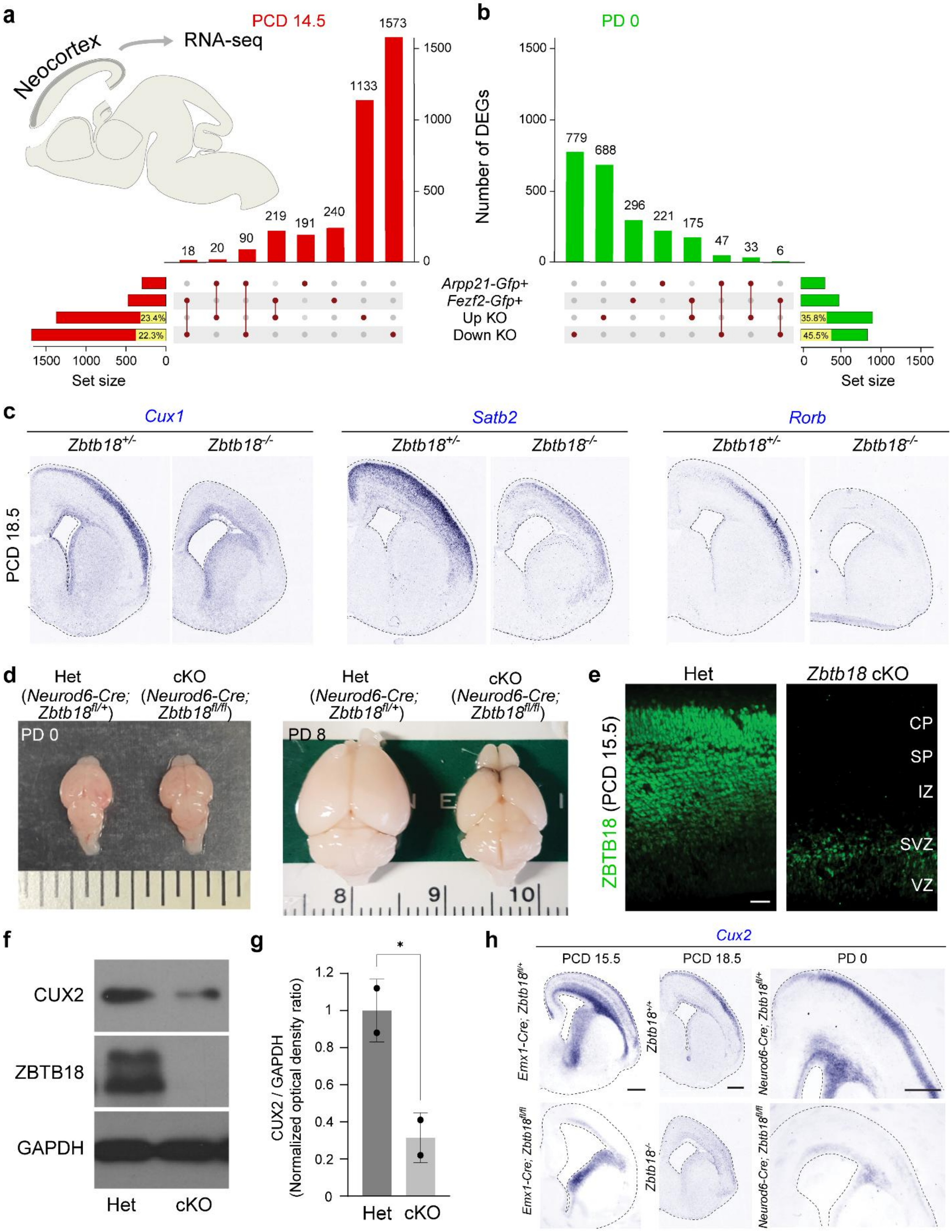
a
PCD 14.5
Neocortex
RNA-seq
18
20
90
219
191
240
1133
1573
Arpp21-Gfp+
Fezf2-Gfp+
Up KO
Down KO
23.4%
22.3%
Set size
b
PD 0
Number of DEGs
779
688
296
221
175
47
33
6
35.8%
45.5%
Set size
c
Cux1
Satb2
Rorb
Zbtb18+/-
Zbtb18-/-
PCD 18.5
d
Het
(Neurod6-Cre; Zbtb18fl/+)
cKO
(Neurod6-Cre; Zbtb18fl/fl)
PD 0
PD 8
e
Het
Zbtb18 cKO
ZBTB18 (PCD 15.5)
CP
SP
IZ
SVZ
VZ
f
CUX2
ZBTB18
GAPDH
Het
cKO
g
CUX2 / GAPDH
(Normalized optical density ratio)
*
Het
cKO
h
Cux2
PCD 15.5
PCD 18.5
PD 0
Emx1-Cre; Zbtb18fl/+
Zbtb18+/+
Neurod6-Cre; Zbtb18fl/+
Emx1-Cre; Zbtb18fl/fl
Zbtb18-/-
Neurod6-Cre; Zbtb18fl/fl

**Extended Data Figure 6. Downregulation of essential IT neuron marker genes in mice lacking ZBTB18**

**a-b**) Bulk RNA-Seq of PCD 14.5 and PD 0 neocortex was performed in whole body *Zbtb18* -/- knockout (KO) and *Zbtb18* +/- (Control) mice and in *Neurod6-Cre*; *Zbtb18*$^{fl/fl}$ *(Neurod6-Cre*; *Zbtb18* cKO) and *Neurod6-Cre*; *Zbtb18*$^{fl/+}$ (*Neurod6-Cre*; *Zbtb18* cHET) (Control) respectively. 82.3% of genes upregulated in *Arpp21-Gfp*+ cells were downregulated in the *Neurod6-Cre*; *Zbtb18* cKO [93 of 113], while 92.4% of genes enriched in *Fezf2-Gfp*+ cells were upregulated in the cKO [219 of 237]. **c**) *In situ* hybridization shows mRNA expression of upper layer marker genes *Cux1, Satb2,* and *Rorb*. Scale bar: 150 µm. **d**) The size of the dorsal brain of *Neurod6-Cre*; *Zbtb18*$^{fl/fl}$ (cKO) mice don't change at PD 0 but appears to be smaller at PD 8 as compared with *Neurod6-Cre;* Zbtb*18*$^{fl/+}$; mice. (e) Immunofluorescence staining of ZBTB18 is shown in the PCD 15.5 Neurod6-Cre; Zbtb18 cKO brain, and ZBTB18 expression is lost in postmitotic neurons but not SVZ progenitors. **f**) Western blot shows ZBTB18 protein expression is abolished, and CUX2 expression is significantly diminished in PD 0 *Emx-Cre; Zbtb18* cKO mouse brain. **g**) Quantified signal intensity shows CUX2 expression is reduced by 70% in *Emx-Cre; Zbtb18* cKO mouse brain compared with control. An unpaired t-test was used to compare groups. The graph represents mean ± s.e.m. * P = 0.0465. **h**) *Cux2* mRNA expression is significantly reduced in the neocortical plate but not the striatum of *Zbtb18* knockout mice at multiple developmental periods. 150 µm. CP, cortical plate; IZ, intermediate zone; SP, subplate; SVZ, subventricular zone; VZ, ventricular zone. All immunofluorescent analyses and *in situ* hybridization represent images selected from multiple sections from at least 3 independent biological replicates. Western blotting is representative of 2 independent biological replicates.

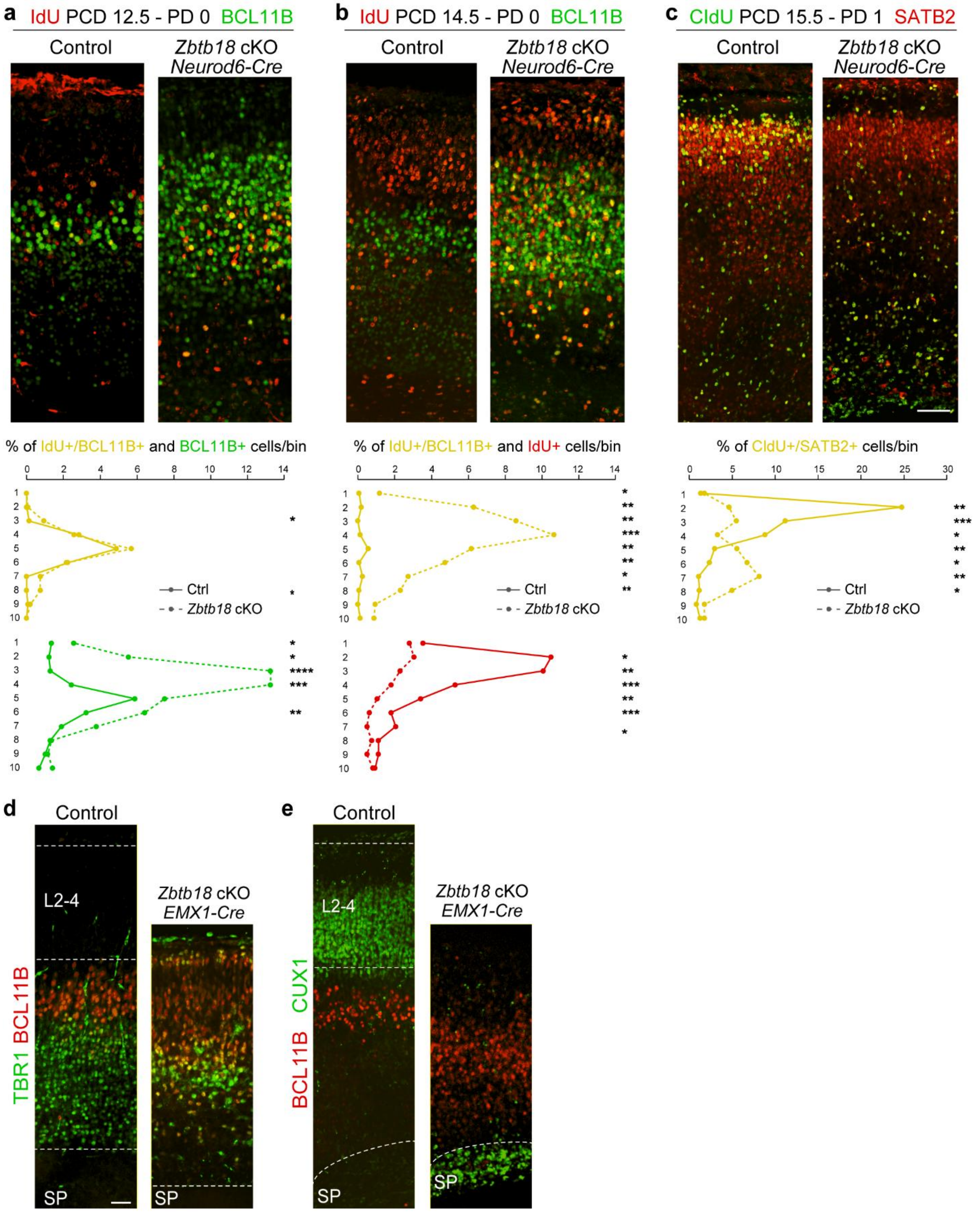
a IdU PCD 12.5 - PD 0 BCL11B
Control
Zbtb18 cKO Neurod6-Cre
% of IdU+/BCL11B+ and BCL11B+ cells/bin
Ctrl
Zbtb18 cKO
b IdU PCD 14.5 - PD 0 BCL11B
Control
Zbtb18 cKO Neurod6-Cre
% of IdU+/BCL11B+ and IdU+ cells/bin
Ctrl
Zbtb18 cKO
c CldU PCD 15.5 - PD 1 SATB2
Control
Zbtb18 cKO Neurod6-Cre
% of CldU+/SATB2+ cells/bin
Ctrl
Zbtb18 cKO
d Control
TBR1 BCL11B
L2-4
SP
Zbtb18 cKO EMX1-Cre
SP
e Control
CUX1 BCL11B
L2-4
SP
Zbtb18 cKO EMX1-Cre
SP

**Extended Data Figure 7. Impact of postmitotic depletion of ZBTB18 on IT neuron development and upper layer formation**

**a)** Top: IdU labeling (red) expected deep layer, BCL11B-immunopositive neurons at PCD 12.5 and immunofluorescence analysis with BCL11B (green) at PD 0 shows a greater number and broader distribution of BCL11B- positive neurons without a concomitant change in BCL11B and, IdU double-positive neurons. Bottom: Quantification of cells per laminar position (bins). **b**) Similar to A, with IdU labeling putative upper layer neurons at PCD 14.5, we observed a significant increase in the number of BCL11B and IdU double-positive neurons at PD 0. **c**) Similar to a and b, with CIdU labeling (green) expected upper layer neurons at E15.5 and immunofluorescence analysis with SATB2 (red) indicating upper layer neurons at P1. *Zbtb18* deficient cells exhibited an alteration in the distribution of SATB2-immunopositive neurons that are also CIdU- positive relative to control mice. For each experiment, 145 to 356 cells from independent sections from at least three different mice were analyzed. T-tests were used to compare control and knockout cell counts per bin. The graph represents mean ± s.e.m. * Bonferroni corrected P-value < 0.05. Please refer to the supplementary table 1 for detailed p values. Scale bar: 100µm. **d-e**) In early postnatal *Emx1-Cre*; *Zbtb18*$^{fl/fl}$ mice brain, layers 2-4 are absent, BCL11B- positive cells populate the top layers of the neocortex, CUX1 expression is absent from upper layers, but present in SP. Scale bars: 50µm. All analyses depicted in this figure were conducted on at least 3 independent biological replicates.

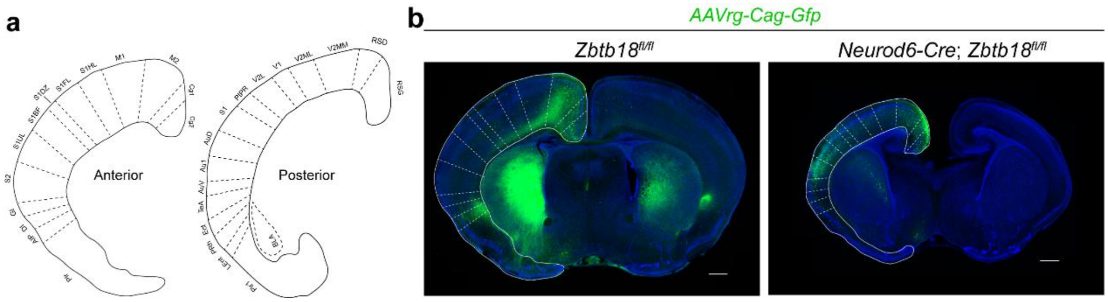


**Extended Data Figure 8. ZBTB18 depletion leads to increased intrahemispheric cortico-cortical association projections originating from the temporal cortex and extending into the mPFC.**

**a**) Coordinates (33) of bins used to analyze tracing experiments depicted in b. **b**) Coronal sections of traced PD 7 brains showing more intensive GFP labeling in GI, S2, S1Ul in *Neurod6-Cre*/+; *Zbtb18fl*fl cKO mice than control. GI, granular insular cortex; S2, secondary somatosensory cortex; S1UL, primary somatosensory cortex, upper lip. Scale bar:

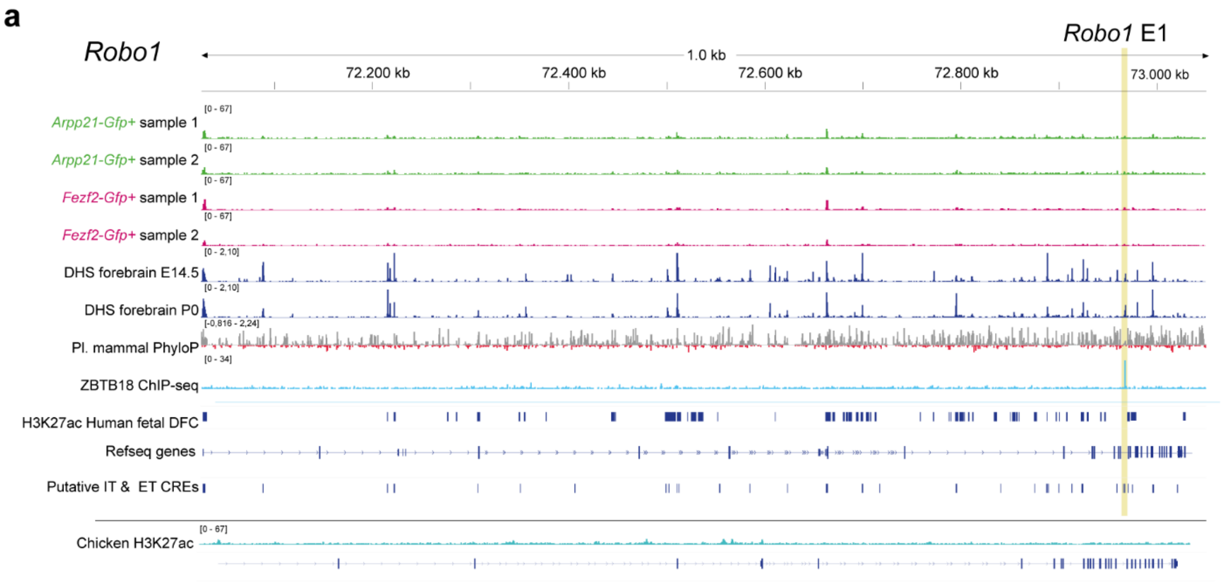


**Extended Data Figure 9**. **ZBTB18 directly binds to the mammalian-specific enhancer *Robo1* E1.**

**a**) Line graphs showing the H3K27ac peaks from, *Arpp21-Gfp*+ cells (line 1 and 2), *Fezf2-Gfp*+ cells (line 3 and 4), from PCD 14.5, and PD 0 forebrain DNaseI Hypersensitivity tracks (DHS) by Digital DNaseI from ENCODE, (line 5 and 6, respectively) and mammal conservation as reference (line 7), ZBTB18-HA ChIP-seq peaks from mouse (line 8), H3K27ac peaks from fetal dlPFC (line 9), gene locus in mice (line 10) H3K27ac peaks collapsed together in the *Arpp21-Gfp*+ IT neurons and *Fezf2-Gfp*+ ET neurons (line 11) and H3K27ac peaks from E17 chicken dorsal pallium (12) at the genomic locus of *Robo1*. The regions detected by H3K27ac and ZBTB18- ChIP seq are in light beige color.

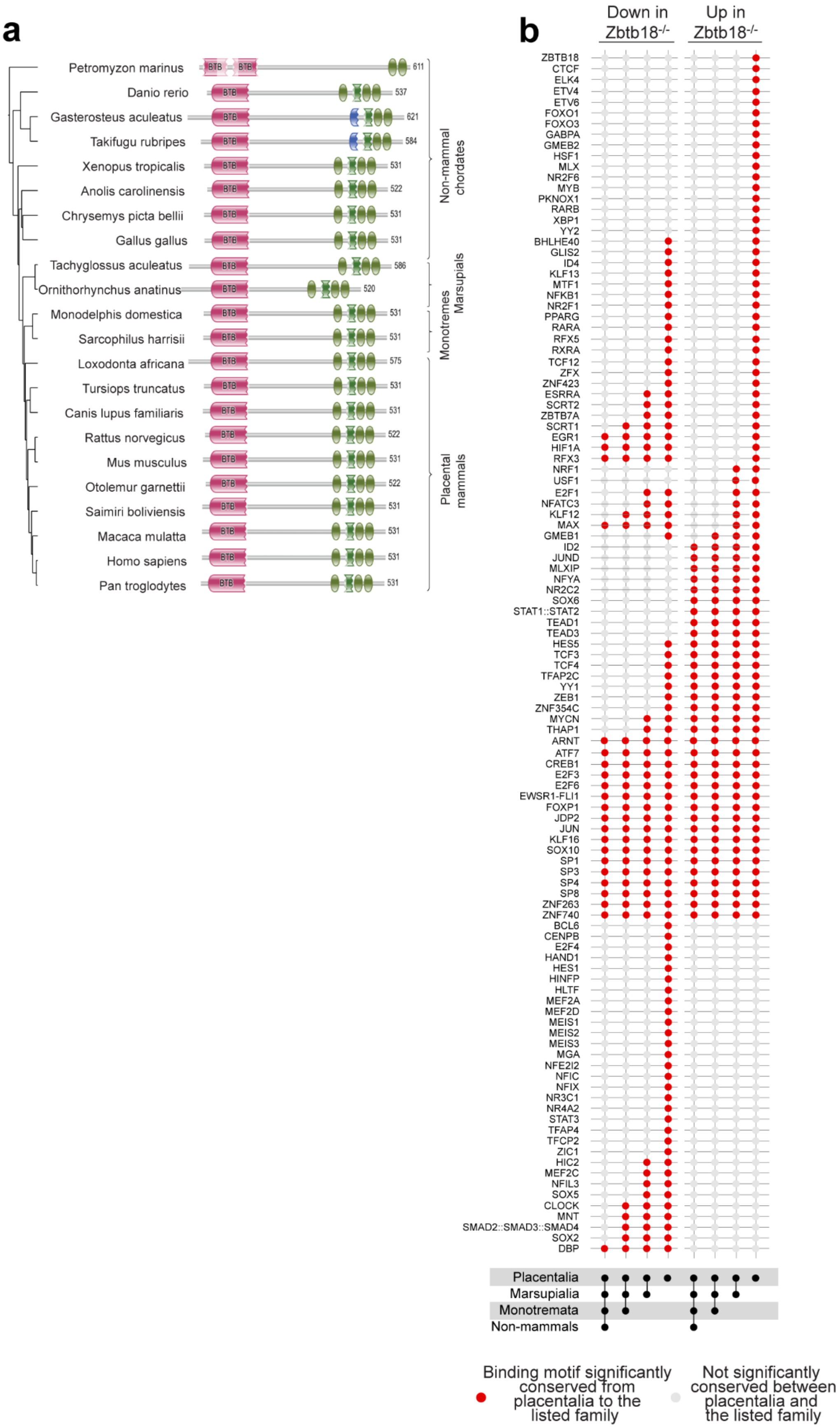
a
Petromyzon marinus
Danio rerio
Gasterosteus aculeatus
Takifugu rubripes
Xenopus tropicalis
Anolis carolinensis
Chrysemys picta bellii
Gallus gallus
Tachyglossus aculeatus
Ornithorhynchus anatinus
Monodelphis domestica
Sarcophilus harrisii
Loxodonta africana
Tursiops truncatus
Canis lupus familiaris
Rattus norvegicus
Mus musculus
Otolemur garnettii
Saimiri boliviensis
Macaca mulatta
Homo sapiens
Pan troglodytes
Non-mammal chordates
Marsupials
Monotremes
Placental mammals
b
Down in Zbtb18-/-
Up in Zbtb18-/-
Placentalia
Marsupialia
Monotremata
Non-mammals
Binding motif significantly conserved from placentalia to the listed family
Not significantly conserved between placentalia and the listed family

**Extended Data Figure 10. ZBTB18 binding motifs exhibit high conservation in putative CREs of mammalian IT neurons.**

**a**) Taxonomic tree of the 23 Chordate species ZBTB18 SST protein sequence features of ZBTB18 were examined. For every species, the Pfam architecture of protein domains is indicated to the right. **b**) The ZBTB18 binding motif, as well as 16 other motifs from the JASPAR database, have consensus sequences enriched in regulatory regions related to genes downregulated in the *Zbtb18* KO that are specifically conserved in placental mammals. Fisher's exact test was used to test for an enrichment of conservation. Red dots represent motifs with a False Discovery Rate corrected P-value < 0.05.

### Extended Data Tables

**Extended Data Table 1: Differential gene expression analysis between *Fezf2-Gfp*+ and *Arpp21-Gfp*+ neocortical cells**

**Extended Data Table 2: H3K27ac peaks in *Fezf2-Gfp*+ and *Arpp21-Gfp*+ neocortical cells analysis**

**Extended Data Table 3: Differentially enriched H3K27ac peaks between *Fezf2-Gfp*+ and *Arpp21-Gfp*+ neocortical cells**

**Extended Data Table 4: Top gene ontology terms of TFs with binding motifs in shared and biased *Fezf2-Gfp*+ and *Arpp21-Gfp*+ H3K27ac peaks that are absent from embryonic chicken dorsal pallium**

**Extended Data Table 5: Gene ontology terms associated with the TFs with binding sites enriched in *Arpp21-Gfp*+ -Biased H3K27ac peaks**

**Extended Data Table 6: Gene ontology terms associated with the TFs with binding sites enriched in *Fezf2-Gfp*+ -biased H3K27ac peaks**

**Extended Data Table 7: Gene ontology terms associated with the TFs with binding sites enriched in shared *Fezf2-Gfp*+ and *Arpp21-Gfp*+ H3K27ac peaks**

**Extended Data Table 8: Enrichment of TF binding motifs in *Arpp21*-Gfp+ biased H3K27ac peaks and H3K27ac peaks related to *Arpp21-Gfp*+ -biased genes**

**Extended Data Table 9: Enrichment of TF binding motifs in *Fezf2*-Gfp+ biased H3K27ac peaks and H3K27ac peaks related to *Fezf2-Gfp*+ -biased genes**

**Extended Data Table 10: Coordinates of the H3K27ac peaks derived from embryonic chicken dorsal pallium in galGal6 and mm10**

**Extended Data Table 11: Coordinates of the ZBTB18 ChIP-Seq peaks identified using four different approaches and their Intersection with H3K27ac datasets from mice and chicken**

**Extended Data Table 12: Differential gene expression in the neocortex of *Zbtb18* knockout Mice compared to heterozygous controls at PCD 14.5**

**Extended Data Table 13: Differential gene expression in the neocortex of *Neurod6-Cre*; *Zbtb18* fl/fl knockout mice Compared to heterozygous controls at PD 0**

**Extended Data Table 14: List of antibodies used in the study**

**Extended Data Table 15: List of the primers used in the study**

### Materials and Methods

#### Mice

All experiments using animals were approved by the Yale University Institutional Animal Care and Use Committee (IACUC) and conducted in compliance with all relevant university, state, and federal guidelines. The day of vaginal plug detection was designated as post-conception day (PCD) 0.5. The day of birth was designated as postnatal day (PD) 0. *Fezf2-Gfp* (MMRRC Cat# 000293-UNC) and *Arpp21-Gfp* (MMRRC Cat# 011848-UCD) transgenic mice were obtained from the gene expression nervous system atlas (GENSAT) [51]. *Cux2-E1-Gfp, Cux2-E2-Gfp*, and *Cux2-E3-Gfp* transgenic mice were generated by delivering linearized DNA constructs by pronuclear injection. 3-7 founders from each line were examined for reproducible GFP expression. *Zbtb18* knockout mice were generated by the Masai lab [13], and *Zbtb18*$^{fl/fl}$ mice were generated by the Heng lab. *Emx1-Cre* (JAX Stock No: 005628) mice, *Cux2-Cre* mice, *Neurod6-Cre* (aka *Nex1-Cre*) mice and *CAG-Cat-Gfp* (IMSR Cat# JAX:024636) mice were previously generated and described [52-55] (Cre Driver Network at NIH Blueprint for Neuroscience Research). Genotyping primers are shown in the Extended Data Table 15.

Mice were provided food and water ad libitum, a 12 hour light; 12 hours dark cycle, veterinary care as provided by the Yale Animal Resource Center, and clean group housing. All mice used for these experiments or bred for these experiments were in good, healthy condition, as approved by the Yale Animal Resource Center and confirmed through regular veterinary monitoring. To maintain genetic diversity, multiple concurrent breeding pairs were maintained, and siblings were never mated. Both males and females were used randomly throughout this study.

#### Tissue preparation and fluorescence-activated cell sorting (FACS)

Neocortices from PD 0.5 *Fezf2-Gfp*, having GFP-expressing neurons enriched in the deep layer (L5-6) predominantly ET neurons (7), and Arpp21-Gfp, having GFP-expressing neurons enriched in predominantly IT neurons [51] mice were dissected under a dissection microscope and minced with a sterile blade. Tails were collected for determining sex and genotyping. Single cell suspensions from neocortical tissue were prepared by dissociation with a papain-based solution [56] and incubated at 37ºC for 15 minutes with intermittent trituration by autoclaved, fire-polished glass pasteur-pipettes. Cells were then pelleted by centrifugation at 4°C for 5 minutes, washed with sterile 1X phosphate buffer saline (PBS), and filtered through a 40μm strainer. Cells were then sorted to collect GFP$^+$ cells by FACS using FACSArial II (BD Biosciences) sorter or Beckman Coulter MoFlo (Beckman) sorter. Hibernate solution (Gibco) supplemented with 2% fetal bovine serum (FBS) (Gibco) was used to collect FACS sorted cells. Cell preparations were maintained at 4°C during the entire process. For RNA-Seq, immediately after FACS, cells were pelleted by centrifugation at 350g at 4°C for 10 min, washed with PBS, pelleted and flash-frozen by liquid nitrogen and stored at -80ºC. For chromatin immunoprecipitation (ChIP) assays with DNA sequencing (ChIP-Seq), FACS sorted GFP$^+$ cells were pelleted by centrifugation at 350g for 10 minutes at 4°C, washed with PBS, and immediately cross-linked with formaldehyde solution at a final concentration of 1% for 10 minutes at room temperature. Glycine (AmericanBio) at final concentration of 125mM was added, and samples were incubated for 5 minutes at room temperature to quench cross-linking. Cells were again washed with PBS and pelleted, then flash-frozen by liquid nitrogen and stored at -80ºC.

### RNA-Seq and initial analysis

Total RNA was extracted from FACS purified cells or neocortical tissue using Trizol reagent per the manufacturer's instructions. Dnase I (Invitrogen) was added to the extracted total RNA and incubated for 15 minutes at 37°C to eliminate DNA contaminants, then subsequently inactivated per the manufacturer's instructions. RNA concentration and integrity were measured using a Nanodrop (ThermoFisher) and TapeStation 2200 (Agilent). Samples with RNA integrity number (RIN) ≥8 were used for subsequent experiments. Libraries were prepared using the TruSeq Stranded Ribozero total RNA preparation kit (Illumina) per the manufacturer's instructions. Libraries were quality-controlled by TapeStation 2200 (Agilent) and sequenced on the HiSeq 2000 platform (Illumina) at the Yale Center for Genome Analysis (YCGA) to generate 75 bp size single-end reads. Sequencing data were quality-controlled by FastQC and aligned to the mouse genome (NCBI38/mm10) using TopHat (v.1.0.13) with up to two mismatches [57]. An average of 40 million uniquely mapped reads were obtained for each sample. Differential expression analysis was performed by the R package DESeq, and principal component analysis (PCA) analysis was performed by the R package prompt. Differential expression of transcripts was detected using a false discovery rate (FDR) <0.01.

### ChIP-Seq and initial analysis

Pooled GFP$^+$ cells from FACS were used for ChIP-Seq. 2.5x $10^7$ cells per condition were cross-linked with a formaldehyde solution (Sigma-Aldrich) at a final concentration of 1% for 10 minutes at room temperature. L-Glycine (AmericanBio) at a final concentration of 125mM was added and incubated for 5 minutes at room temperature to quench the cross-linking. Cells were washed by PBS, then disrupted by lysis buffer I (50 mM HEPES-KOH pH 7.5, 140 mM NaCl, 1M EDTA pH 8.0, 10% Glycerol, 0.5% NP-40, 0.25% TritonX-100, 1X Protease inhibitor) for 20 minutes at 4°C and lysis buffer II (200 mM NaCl, 1M EDTA pH 8.0, 0.5 mM EGTA pH 8.0, 10 mM Tris-HCl pH 8.0, 1X Protease inhibitor) for 10 minutes at room temperature. Cells were centrifuged at 300g for 15 minutes at 4°C and pellets were dissolved in 400-600µl lysis buffer III (1mM EDTA pH 8.0, 0.5mM EGTA pH 8.0, 10 mM Tris-HCl pH 8.0, 0.5% Sarkosyl, 1X Protease inhibitor) prior to being sheared into 200-500 bp size fragments with a sonicator (Bioruptor, Diagenode). Dyna beads protein G (Invitrogen) was pre-blocked with 5mg/ml ice-cold bovine serum albumin (BSA) and incubated with 5µg anti-H3K27ac antibody (Abcam) at 4°C with constant rotation for 12 hours. 25µg chromatin was added to the bead-antibody complex mixture per reaction and incubated with constant rotation for 16 hours at 4°C. Beads were washed with ice-cold radio-immuno-precipitation assay (RIPA) buffer (Thermo Scientific) 8 times, rinsed by 1XTris-EDTA (TE) solution, eluted by adding 200µl ChIP elution buffer (1% SDS, 1XTE), and incubated in a shaker for 20 minutes at 65°C. ChIP DNA was incubated for 12 hours at 65°C for reverse cross-linking, and subjected to RNAse A (Thermo Scientific) treatment (1 hour, 37°C) and Proteinase K (Sigma-Aldrich) treatment (2 hours, 55°C), then purified on polymerase chain reaction (PCR) purification columns. For input control, 5µg chromatin from whole-cell-extract from each sample was also treated by reverse cross-linking, RNAse A (Thermo Scientific, EN0531), and Proteinase K (Sigma-Aldrich, 3115887001) together with IP samples and purified by PCR purification columns. DNA amounts were quantified by PicoGreen assay (Thermo Scientific, P7589). 5ng IP DNA and input from each sample were used to prepare ChIP libraries with Truseq ChIP library preparation kit (Illumina, IP-202-1012) per manufacturer's instructions. Libraries were size selected to enrich

300-400 bp size fragments and quality controlled and sequenced on Hiseq 2000 platform (Illumina) (YCGA). Approximately, 20-25 million reads were obtained from each sample. FASTA files were mapped to the mouse genome (NCBI37/mm10) using TopHat v.1.0.13 (http://tophat.cbcb.umd.edu/) and Bowtie 2 (http://bowtie-bio.sourceforge.net/bowtie2/index.shtml)[58]. Peaks were called by MACS2 (SCR_013291) [59].

#### H3K27ac ChIP-Seq in the chicken

Embryonic day (E) 15 or Hamilton-Hamburger stage (HH) 41 [60] chicken eggs were obtained from Charles River Laboratories and incubated in the lab for two days at 37°C in the humidified chamber. The dorsal pallial regions- hyperpallium (H) and mesopallium (M) were micro-dissected from the E 17 (HH 43) chicken embryos. 10 pallia were pooled and cross-linked with a formaldehyde solution (Sigma-Aldrich) at a final concentration of 1% at room temperature for 10 minutes per sample. L-Glycine (AmericanBio, 56-40-6) at final concentration of 125mM was added and incubated for 5 minutes at room temperature to quench the cross-linking. Cells were washed by PBS thrice and cells were lysed in the hypotonic solution (50 mM Tris-Cl pH 7.5, 0.5% NP40, 0.25% Sodium Deoxycholate, 0.1% Sodium Dodecyl Sulfate (SDS), 150 mM NaCl) on ice for 10 minutes to obtain the nuclei. Nuclei were centrifuged at 600g for 5 minutes at 4°C and pellets were resuspended in the SDS lysis buffer (1% SDS, 10mM EDTA and 50mM Tris-Cl, pH 8.1) prior to being sheared into 200-500 bp size fragments using sonicator (M220 Focused-ultrasonicator, Covaris). The sheared DNA was diluted with the ChIP dilution buffer (0.01% SDS, 1.1% Triton X- 100, 1.2mM EDTA, 16.7mM Tris-Cl, pH 8.1, 167mM NaCl) and was precleared with Magnetic Protein A/G Beads (Millipore Sigma) for 1 hour at 4°C. Beads were discarded and anti-H3K27ac antibody (Abcam, ab4729) was added. Samples were incubated on constant rotation for overnight at 4°C. Magnetic Protein A/G Beads (Millipore Sigma) blocked with 1mg/ml BSA (Sigma-Aldrich) and tRNA were added to the chromatin- antibody complexes for 4 hours at 4°C. Beads were washed with low salt (0.1% SDS, 1% Triton X-100, 2 mM EDTA, 20mM Tris-Cl, pH 8.1, 150mM NaCl), high salt (0.1% SDS, 1% Triton X-100, 2mM EDTA, 20mM Tris-Cl, pH 8.1, 500mM NaCl), LiCl (0.25M LiCl, 1% IGEPAL CA630, 1% deoxycholic acid (sodium salt), 1mM EDTA, 10mM Tris-Cl, pH 8.1), and TE (AmericanBio, AB14033-01000), sequentially for 3 mins each. ChIP DNA was incubated overnight at 65°C for reverse cross-linking and subjected to RNAse A (Thermo Scientific, EN0531) treatment (37°C, 1 hour) and Proteinase K (Sigma-Aldrich, 3115887001) treatment (55°C, 2hours), then purified on PCR purification columns. For input control, 5µg cross-linked chromatin from each sample was also treated for reverse cross-linking, RNAse A (Thermo Scientific, EN0531), and Proteinase K (Sigma-Aldrich, 3115887001) together with immunoprecipitated (IP) samples and purified by PCR purification columns. DNA amounts were quantified by PicoGreen assay (Thermo Scientific, P7589). 10ng IP DNA and input from each sample were used to prepare library using KAPA HyperPrep Kit (Roche, KK8500) as per manufacturer's instructions and sample multiplexing was done using CD dual indexes (Illumina) (YCGA). Libraries were size selected to enrich 300-400 bp size fragments and quality controlled and sequenced on Hiseq 2000 platform (Illumina). Approximately, 60 million reads were obtained per sample. Reads were mapped to the chicken genome (Galgal6) using Bowtie2 v2.4.2 (RRID:SCR_016368, http://bowtie-bio.sourceforge.net/bowtie2/index.shtml), (Langmead and Salzberg, 2012). Duplicates were removed and unique reads with MAPQ>20 were selected using SAMtools v1.16 (RRID:SCR_003030, https://github.com/samtools/samtools). Peaks were called using MACS2 v2.2.7.1 (RRID:SCR_008036).

**ZBTB18 ChIP-Seq in the mouse**

Epitope-tagged ChIP were performed to identify ZBTB18 binding sites. Cortices from PD 0 were isolated and were subjected to enzymatic dissociation. Cells from 15 cortices were pooled and treated as one sample. Cells were washed and transfected with HA-tagged Zbtb18 using Amaxa Basic Nucleofector™ Kit (Lonza BioSciences, VPI-1003) for primary mammalian neurons per manufacturer's instructions. After 48 hours of culture, media was removed, and cells were cross-linked with a formaldehyde solution at a final concentration of 1% at room temperature for 10 minutes. L-Glycine (AmericanBio, 56-40-6) at a final concentration of 125mM was added and incubated for 5 minutes at room temperature to quench the cross-linking. The cells were scraped and collected in a 50 ml conical tube and lysed in the hypotonic solution (50 mM Tris-Cl pH 7.5, 0.5% NP40, 0.25% Sodium Deoxycholate, 0.1% SDS, 150 mM NaCl) on ice for 10 minutes to obtain the nuclei. Nuclei were centrifuged at 600g for 5 minutes at 4°C and pellets were resuspended in the SDS lysis buffer (1% SDS, 10mM EDTA and 50mM Tris-Cl, pH 8.1) prior to being sheared into 200-500 bp size fragments using a sonicator (M220 Focused-ultrasonicator, Covaris). The sheared DNA was diluted with the ChIP dilution buffer (0.01% SDS, 1.1% Triton X-100, 1.2mM EDTA, 16.7mM Tris-Cl, pH 8.1, 167mM NaCl) and was precleared with magnetic protein A/G beads (Thermo Scientific) for 1 hour at 4ºC. For epitope-tagged ChIP, 5μg anti-HA antibody (Millipore Sigma, 11867423001) was used. Samples were incubated on constant rotation for overnight at 4ºC. Magnetic protein A/G beads (Millipore Sigma, 88803) were blocked with 1mg/ml BSA (Sigma-Aldrich, A2153) and tRNAs were added to the chromatin-antibody complexes for 4 hours at 4ºC. Beads were washed with low salt (0.1% SDS, 1% Triton X-100, 2 mM EDTA, 20 mM Tris-Cl, pH 8.1, 150 mM NaCl), high salt (0.1% SDS, 1% Triton X-100, 2 mM EDTA, 20 mM Tris-Cl, pH 8.1, 500 mM NaCl), LiCl (0.25 M LiCl, 1% IGEPAL CA630, 1% deoxycholic acid (sodium salt), 1 mM EDTA, 10 mM Tris-Cl, pH 8.1), and TE (AmericanBio), sequentially for 3 minutes each. ChIP DNA was incubated overnight at 65°C for reverse cross-linking and subjected to RNAse A treatment (37°C, 1hour) and Proteinase K treatment (55°C, 2 hours), then purified on PCR purification columns. For input control, 5μg cross-linked chromatin from each sample was also treated by reverse cross-linking, RNAse A (Thermo Scientific), and Proteinase K (Millipore Sigma) together with IP samples and purified by PCR purification columns. DNA amounts were quantified by PicoGreen assay (Thermo Scientific, P7589). 10 ng IP DNA and input from each sample were used to prepare library using KAPA HyperPrep kit (Roche, KK8500) as per manufacturer's instructions and multiplexing using CD dual indexes (Illumina). Libraries were size selected to enrich 300-400 bp size fragments and quality controlled and sequenced on Hiseq 2000 platform (Illumina) (YCGA). Approximately 60 million reads were obtained per sample. Reads were mapped to the chicken genome (mm10) using Bowtie2 v2.4.2. Duplicates were removed and unique reads with MAPQ>20 were selected using SAMtools (v1.16). We explored different options to determine putative Zbtb18 peaks, in terms of replicates and parameters, including using TRANSFAC [61], PROMO [62], MatInspector [63] and JASPAR databases [64]. Hence, peaks were called using 4 different approaches: using MACS2 [59] and its default parameters and requiring the peak to be present in at least two biological replicates, using MACS2 with the "--nomodel" and "--nolambda" parameters and also requiring consistency across replicates, using MACS2 with default parameters but only requiring the peak to be present in replicate 2, and using MACS3 with the peaks of the replicate 2.

**TF binding site prediction of candidate CREs**

We obtained the DNA sequence for all 62,448 H3K27ac peaks using *twoBitToFa* [65] tool (https://genome.ucsc.edu/goldenPath/help/twoBit.html). We then ran a tool called Find Individual Motif Occurrences (FIMO) (https://meme-suite.org/meme/doc/fimo.html), with default parameters to predict TF binding sites in those sequences [66]. We used the JASPAR 2016 Core dataset [67] in meme format containing 635 motifs. For the ET neurons-specific TF binding sites enrichment analysis, we calculated the number of bases present in the binding sites for a particular TF and but not in the binding sites within IT neuron-associated peaks and compared to the same numbers in ET neuron-associated peaks via Fisher's exact test. Motifs with an FDR-corrected p-value < 0.05 were considered significantly enriched.

**Generation of enhancer reporter transgenic mice**

Putative enhancers *Cux2-E1*, *Cux2-E2* and *Cux2-E3* were amplified from mouse genomic-DNA and cloned into pBgn-Gfp vector (the Sestan lab), then sequence verified by Sanger sequencing (YCGA). Enhancers were placed upstream of the human BGN minimal promoter [68] to drive GFP expression. Primers and oligonucleotides used for cloning DNA constructs are shown in the Extended Data Table 15. Enhancer-inserted plasmids were linearized using appropriate restriction enzymes and size-selected by gel electrophoresis, then purified by phenol/chloroform extraction. A final concentration of 2.5ng/μl DNA was used for pronuclear injection. At PD 0, pups were examined for GFP expression under a fluorescence microscope, and tails were collected to genotype and confirm the presence of *Gfp loci*. For GFP+ founders, brains were harvested, and proceeded with immuno-histological analysis to examine the GFP expression pattern. 3-7 founders from each transgenic line with stable GFP expression patterns were acquired and analyzed.

**Quantitative RT-PCR**

Total RNA was extracted using Trizol (Invitrogen, 15596018) reagent from freshly isolated neocortical tissue from PCD 15.5 and PD 0 wildtype, *Neurod6-Cre*; *Zbtb18* $^{fl/fl}$ cKO and *Neurod6-Cre; Zbtb18*$^{fl/+}$ (control) mice and subjected to Dnase I (Invitrogen, AM1907) treatment as previously described. cDNA was synthesized using reverse transcriptase (Invitrogen, 18080093) per the manufacturer's instructions, and quantitative PCR was performed in triplicate for each sample using RT-PCR machine (iQ5 system, BioRad) with primer sets spanning exon junctions of the targeted transcripts. Identical or near-identical sized transcript fragments from the mouse *Tbp* gene were used as an internal control, and the expression level of each gene was normalized to wild-type for relative fold changes. Sequences of primers used are provided in the key-resource table. Unpaired, two-tailed t-test was used to detect differences between samples.

**Single cell-RNA-Seq analysis across species**

To assess the expression patterns of *Zbtb18* across different species, we reanalyzed public single-nucleus transcriptome datasets for amygdala in turtle (*Trachemys scripta elegans*), lizard (*Pogona vitticeps*), chicken (*Gallus gallus*) and mouse (*Mus musculus*). We checked the expression level of *Cux2*, *Satb2*, *Robo1*, *Bcl11b*, *Ccdc1167*, *Rpl38-ps1*, *Arhgef16*, *n-R5s136*,

*3830408C21Rik*, *Gm24089* and *Gm42489* across major cell types, as identified by the original paper [12,28-30]. For cross-species comparison, we only included the excitatory neurons.

**Postmortem human tissue**

De-identified postmortem human brain tissue was acquired from the Sestan lab collection or the the National Institutes of Health (NIH) Neurobiobank (https://neurobiobank.nih.gov), as previously published [69]. All tissue was collected with the informed consent of parental or next-of-kin and the approval of all relevant review boards or committees of the Yale University School of Medicine and the NIH. Similarly, tissues were handled in accordance with the appropriate constraints, regulations, and ethical guidelines for the research use of human brain tissue set forth by the NIH (http://bioethics.od.nih.gov/humantissue.html) and the world medical association declaration of Helsinki (https://www.wma.net/policies-post/wma-declaration-of-helsinki-ethical-principles-for-medical-research-involving-human-subjects/). The sample used in this study were used without regard for gender and analyses into the influence or sex-specific characteristics of male versus female samples were not considered in this study.

Samples were fixed in 4% paraformaldehyde (PFA) (Electron Microscopy Sciences) for 2 days at 4°C. Tissue sections were mounted and dried overnight. Antigen retrieval was performed using R-Buffer A pH 6.0 (Electron Microscopy Sciences) and the benchtop Antigen retriever device (Electron Microscopy Sciences) according to the manufacturer's instructions. Sections were washed in PBS (3 x 15 minutes) and incubated in blocking solution containing 5% (v/v) normal donkey serum (Jackson ImmunoResearch Laboratories), 1% (w/v) BSA, and 0.4% (v/v) Triton X-100 (Sigma-Aldrich) in PBS for 1 hour at room temperature. Primary antibodies were diluted ZBTB18 (Proteintech, 12714-1-AP), 1:1000; BCL11B (Abcam, ab18465), 1:500; SATB2 (GenWay Biotech Inc., GWB-9F2D9F), 1:200) in blocking solution and incubated with tissues sections for two nights at 4°C. Sections were washed with PBS (3 x 15 minutes) prior to being incubated with the appropriate fluorescent secondary antibodies (Jackson ImmunoResearch Labs) for 1.5 hours at room temperature. All secondary antibodies were raised in donkey and diluted at 1:250 in blocking solution. Finally, sections were washed in PBS with 0.3% Triton X-100 and treated with the Autofluorescence Eliminator Reagent (Millipore Sigma, 2160) according to manufacturer's instructions, and cover-slipped with aqueous mounting media (Vectashield, Vector laboratories). Sections were digitized using confocal microscope (Zeiss, LSM 510 meta) and images were assembled in Zeiss Zen, ImageJ, Adobe Photoshop, and Adobe Illustrator.

**Cell line and transfection**

Neuro-2a cells (ATCC, CCL-131) were maintained in DMEM medium (Gibco) supplied with 10% FBS (Gibco), L-Glutamine (Thermofisher Scientific), Penicillin (Gibco) and Streptomycin (Gibco). Cells were passaged every 2-3 days upon reaching 80% confluency. For transfection experiments, Neuro-2a cells were seeded at an appropriate density the day before transfection, and when the cells reached 60-70% confluency, transfection was performed 20-24 hours later, with Lipofectamine 2000 (Thermo Fisher Scientific, 11668027) mixed with appropriate vectors at a DNA: Lipid ratio of 1:3. Either 24 or 48 hours post-transfection, cells were dissociated with 0.25% trypsin (Gibco), washed with PBS, and processed for corresponding assays.

**Primary neuronal culture, AAV infection and immunostaining**

Neocortices from PD 0.5 *Zbtb18*$^{fl/fl}$ and *Zbtb18*$^{fl/+}$ mice were isolated and dissociated using a papain-based dissociation solution (Worthington Biochemical Corporation). Briefly, mice were decapitated, and brains were quickly isolated, meninges were peeled-off with fine-tip tweezers, and neocortex from each mouse was dissected under a dissection microscope, minced with a sterile blade, and collected into a 15 ml falcon tube with 1ml dissociation solution. Complete dissociation was achieved by incubation for 10 minutes at 37°C with intermittent trituration by a fire-polished glass pasteur-pipette (3-5 times). Cell suspensions were centrifuged at 300g for 3 minutes at room temperature, then washed with sterile PBS and filtered through a 40μm strainer. 50,000 neurons were plated into each cell culture chamber. Neurobasal medium (Gibco), supplemented with 2% B-27 supplement (Thermo Scientific), 2mM L-Glutamine (AmericanBio), and 1 M Glucose (AmericanBio), was then added. Then 12 hours after plating, 2 μl of AAV-CAG-Cre and AAV-CAG-Gfp viral-particle suspension was added to infect cells for 24 hours, then media was exchanged with fresh culture medium. Cultures were incubated at 37°C in a 95% air/5% $CO_2$ humidified incubator for 10 days prior to fixation and imaging. Alternatively, we replicated these experiments using 1x $10^{12}$ GC/ml titers of AAV-Synapsin-Cre-GFP (AAV Serotype 8; SignaGen Laboratories, SL100883). Cells were fixed in 4% paraformaldehyde (Sigma-Aldrich) at room temperature on day 10 and immunostaining was subsequently performed. Briefly, cells were washed twice with cold PBS solution, fixed with 4% formaldehyde for 10 minutes, and then treated with 0.5% Triton-X100 in PBS solution for 15 minutes. After blocking with 5% BSA in PBS solution for 1 hour, cells were incubated with primary antibodies at 4°C overnight. Secondary antibodies were subsequently incubated for 1 hour at room temperature on the next day.

***In utero* electroporation**

*In utero* electroporation was performed on PCD 14.5 and PCD 15.5 timed-pregnancy embryos (n=3-6 for each condition) and littermates were used as controls. 0.5μl DNA preparation (4μg/μl DNA mixed with 0.05% Fast Green FCF dye (Sigma-Aldrich) was injected into the lateral ventricle of the embryos and electroporated using square-wave pulse electroporator (Harvard Apparatus, Inc., BTX) at 35V-38V, 5 pulses, 50ms ON and 950ms OFF, to deliver DNA constructs to the ventricular zone. At PD0, pups were screened for GFP or RFP expression under a fluorescence-attached dissection microscope. Pups with the fluorescence signal were euthanized, and brains were analyzed as previously described.

**Neuronal tracing and imaging**

Neonatal mice injections were performed with a motorized micro-injector (RWD system; cat# MM-500 and, R480). Briefly, mice at PD 3 were anesthetized through hypothermia and ~50nl retrograde AAV viral particles carrying pCAG-Gfp cassette (Addgene, 37825-AAVrg, titer ≥ $7x10^{12}$vg/ml) were injected into medial-prefrontal cortex (mPFC) visually with the guide of stereo binocular microscope. At PD 7 the injected mice that were properly targeted were perfused with ice-cold PBS followed by 4% PFA (Electron Microscopy Sciences). The collected brains underwent post fixation at 4°C overnight and washed with PBS before cutting coronally at 60μm thickness. The sections were stained with anti-GFP antibody (Abcam, ab13970) before imaging. Images were taken using an automated slide scanner (Olympus, VS200) with 20x objective. For analysis, the images from wildtype mice were aligned with the Allen brain atlas coordinates at PD 6 and the *Neurod6-Cre Zbtb18* cKO was approximated with wildtype counterpart. Cell numbers from each bin were counted using ImageJ tool (RRID:SCR_003070). Total cell number from the neocortex as shown in C from 3 mice of each genotype was compared. For the percentage of distribution, we divide the cell number from each bin by the total number of cells in the neocortex and the comparisons between each bin were made with unpaired t test.

**BrdU/CldU/IdU birth-date labeling**

5′-bromo-2′-deoxyuridine (BrdU) (Sigma-Aldrich, B5002) or 5-Chloro-2-deoxyuridine (CidU) (Sigma-Aldrich, C6891) were dissolved in distilled water at 100mg/ml, and 5-Iodo-2-deoxyuridine (IdU) (Sigma-Aldrich, I7125) was dissolved in distilled water at 50mg/ml and stored at -20°C in the dark. At the proscribed ages, a single-dose of IdU, BrdU, or CidU was re-dissolved by intermittent vortexing and given to the timed-pregnant mice by intra-peritoneal injection at a dosage of 1mg/20g bodyweight (10uL BrdU/CidU or 20uL IdU). Pups were euthanized within the PD 1 and brains were fixed in 4% PFA (Electron Microscopy Sciences) for 12 hours at $4^{o}$ C. Heterozygous littermates were used as controls. Brains were embedded with 4% agarose and sectioned at 50µm thickness on a vibratome (Leica, VT1000S). Sections were treated with 2M HCL at room temperature for 30 minutes and co-immunostained with BrdU (Sigma-Aldrich) as well as other markers for immunofluorescence.

**Recombinant DNA**

DNA constructs used for making transgenic mice were sub-cloned from the pBgn-GFP (Sestan lab) backbone with *Cux2-E1*, *Cux2-E2* and *Cux2-E3* inserted. Vectors for over-expression experiments were sub-cloned from *pCAGIG*-backbone (Addgene, 11159) with inserts of the full coding sequences (CDS) of the following genes: mouse *Zbtb18* (aka *Rp58, Znf238, Zfp238)*, BC054529; mouse *Pou3f2*, NM_008899; mouse *Trim28* (aka *Rnf96)*, BC058391; mouse *Lhx2*, BC055741; mouse *Satb2*, BC138626; mouse *Hdac2*, BC138517; mouse *E2F1*, BC052160; mouse *Sin3A*, BC052716; mouse *Foxn2*, NM_001355743; mouse *Coup-tf1* (aka *Nr2f1*), BC108408, mouse *Robo1*, NM_019413.2. CDS in DNA constructs used for IUE experiments were cloned into an expression construct with a *Neurod1*-promoter [70] and verified by Sanger sequencing (YCGA). Other DNA constructs used in this manuscript include *pCAG-Cre, pCALNL-GFP, pCAGIG, pCALSL-RFP, pNeurod1-GFP* and subcloned plasmids. *pCAG-Cre, pCALNL-GFP, pCAGIG* were gifts from Connie Cepko (Addgene plasmid #11159, Addgene plasmid #13770, Addgene plasmid #13775), and *pCALSL-RFP* and *pNeurod1-GFP* were previously described [70,71]. Vectors carrying enhancers for luciferase assay were sub-cloned from pGL4.24 backbone (Promega). For enhancer mutagenesis, desired sites within specific enhancers were mutated using a site-directed mutagenesis kit (New England Biolabs, Q5 Site-Directed Mutagenesis Kit) per the manufacturer's instructions. Mutated products were validated by Sanger sequencing. Oligos and primers used for cloning are in Extended Data Table 15.

### Luciferase assay

Neuro-2a (ATCC, CCL-131) cells were plated into 96-, or 24-well plates at a density of 10,000, or 50,000 cells per well, respectively. Sixteen hours post-plating, cells were transfected with a DNA mixture of 100 ng and 500 ng over-expression plasmids, 30 ng and 500 ng firefly-luciferase (pGL4.24; Promega) plasmids containing candidate enhancers, together with 20 ng and 25ng pRL-SV40 plasmid (Promega) as a control for 96- and 24-well plates, respectively with the assistance of lipofectamine 2000 (Thermo Fisher Scientific). Transfected cells were lysed and assayed 24-48 hours post-transfection using the Dual-Luciferase Reporter Assay System (Promega) per the manufacturer's instructions. Relative luciferase activity was calculated by normalizing the firefly-luciferase luminescence to the renilla-luciferase luminescence value. Luciferase activity was measured and quantified by GloMax-Multi Detection System (Promega). Primers used for amplifying the candidate enhancer and promoter regions are shown in the Extended Data Table 15. Experiments were performed in triplicate or quadruplicate and p-values were calculated using unpaired two-tailed t-tests to compare the over-expression and control. To perform the luciferase assay on multiple mutated enhancers, a one-way ANOVA test was used to detect differences among more than two enhancers. Alpha value=0.05.

### *In situ* hybridization for the human, mouse and chicken brain sections

RNA probes for mouse and human were generated using respective neocortical tissues cDNA as template (mouse *Zbtb18*, ENSMUSG00000063659; mouse *Cux2*, ENSMUSG00000042589; mouse *Satb2*, ENSMUSG00000038331; mouse *Robo1*, ENSMUSG00000022883; mouse *Robo2*, ENSMUSG00000052516; mouse *Robo3*, ENSMUSG00000032128; mouse *Rorb*, ENSMUSG00000036192; mouse *Dcc*, ENSMUSG00000060534; human *ZBTB18*, ENSG00000179456; human *CUX2*, ENSG00000111249) by TA-cloning kit (Invitrogen) and followed by *in vitro* transcription (Roche) per the manufacturer's instructions. For chicken probes, the cDNA fragments coding for Satb2 (NM_001199110), and Cux2 (XM_415167), were amplified by reverse transcriptase-polymerase chain reaction (RT-PCR) and subcloned into the plasmid vector pTA2 (Toyobo).Templates purified by phenol/chloroform extraction and Digoxigenin-labelled probes were synthetized using T3 (Roche) and T7 RNA polymerases (Roche) respectively, and RNA labelling mix (Roche) according to the manufacturer's instructions. Probes were purified by phenol/chloroform extraction, quantified, quality controlled, and stored at -80°C until hybridization. Primers used for generating probes are provided in the Extended Data Table 15.

For single-color *in situ* hybridization (ISH), free-floating or slide-mounted cryo-sections at 20-30µm thickness were processed as previously described protocol [72]. Briefly, brains were fixed overnight at 4°C in 4% PFA (Electron Microscopy Sciences) diluted in DPBS (Thermo Fisher Scientific), equilibrated for 12 hours at 4°C in 10% sucrose, and another 12 hours at 4°C in 30% Sucrose (AmericanBio) in DPBS (Thermo Fisher Scientific). Fixed brains were then embedded in OCT (Scigen) and sliced on a cryostat (Leica Biosystem, CM1800). Slides were stored at -80°C until processed for *in situ* hybridization. Sections were first post-fixed in 4% PFA (Electron Microscopy Sciences) in PBS for 15 minutes at room temperature, washed with PBS, and submerged in hybridization buffer (5X SSC, 50% Formamide, 5X Denhardt's solution, 500 µg/ml of salmon sperm DNA and 250 µg/ml of Torula yeast RNA) supplemented with 1000ng/ml of the appropriate digoxigenin-labeled probe at 70°C overnight. Sections were washed two times for 60 minutes at 70°C in 2X SSC, 50% formamide, 0.1% Tween, washed in 100 mM Tris-Cl pH7.5, 150 mM NaCl, 0.1% Tween, blocked with 10% sheep inactivated serum (Sigma-Aldrich) and incubated overnight at 4°C with an anti-digoxigenin antibody conjugated to alkaline phosphatase (1:2000, Roche). Sections were then rinsed in the substrate buffer (100mM Tris-Cl pH9.5, 100mM NaCl, 50mM $MgCl_2$, 0.1% Tween-20) before being overlaid with NBT/BCIP substrate (Roche). Revelation was done at room temperature in the dark until the desired signal was reached. Finally, sections were rinsed in DPBS (Thermo Fisher Scientific), post-fixed with 4 % PFA (Electron Microscopy Sciences) in DPBS (Thermo Fisher Scientific), washed in water and mounted with permount mounting medium (Electron Microscopy Sciences).

For two color ISH, probes were synthesized by either DIG-labeled (Roche) or fluorescein-labeled (Roche) RNA labeling mixes *in vitro* transcription (Roche); free-floating or slide-mounted cryo-sections at 20-30µm thickness were post-fixed in 4%PFA (Electron Microscopy Sciences) for 15 minutes, washed, and hybridized overnight at 70ºC in 14ml hybridization solution with 500ng/ml digoxigenin (DIG)-labeled (Roche) human *ZBTB18* probe and 500ng/ml fluorescein-labeled (Roche) human *CUX2* probe for human tissue. For mouse tissue, 500ng/ml digoxigenin (DIG)-labeled mouse *Zbtb18* and 500ng/µl fluorescein-labeled mouse *Cux2* probes were used. The signals were sequentially detected with an alkaline phosphatase-conjugated anti-DIG antibody (Millipore-Sigma) and NBT/BCIP substrate (Roche), and horseradish peroxidase conjugated anti-fluorescein antibody and Horseradish peroxidase activity was amplified firstly with TSA (AKOYA) then further strengthen with 3,3′-Diaminobenzidine tetrahydrochloride hydrate (DAB) (Sigma-Aldrich).

**Immunostaining and immunoblotting**

Brains dissected from embryonic and neonatal mice were fixed by immersion in 4% PFA (Electron Microscopy Sciences) overnight at 4°C. Adult brains were perfused by 10 ml PBS and then 10 ml 4% PFA (Electron Microscopy Sciences), then isolated and post-fixed by immersion in 4% PFA (Electron Microscopy Sciences) overnight at 4°C. Brains were sectioned at a thickness of 50µm (postnatal) and 80 µm (embryonic) on a vibratome (Leica, VT1000S). Brain sections were blocked using blocking solution (5% BSA, 10% donkey serum, 0.3% Triton X-100 in 1xPBS solution) for 1 hour at room temperature and incubated with appropriate primary antibodies for 12-24 hours at 4°C, washed with 1xPBS 3 times, and incubated with appropriate secondary antibodies for 1 hour at room temperature. DAPI was used to stain nuclei. Antibody dilutions used for immunostaining were as follows: anti-ZBTB18 (rabbit, 1:1000, Protein Tech), anti-L1CAM (rat, 1:300; Millipore), anti-GFP (chicken, 1:3,000; Abcam), anti-RFP (Rabbit, 1:1,000, Abcam), anti-CUX1 (rabbit, 1:250; Santa Cruz), anti-CUX2 (rabbit, 1:250, Abcam), anti-SATB2 (mouse, 1:200; Genway), anti-BCL11B (rat, 1:500; Abcam), anti-TBR1 (rabbit, 1:250; Santa Cruz), anti-TBR2 (mouse,1:1000, Invitrogen), anti-LHX6 (mouse,1:300, Santa Cruz) anti-SOX2 (goat,1:250; R&D Systems), anti-LHX2 (goat, 1:250; Santa Cruz), anti-LHX6 (goat, 1:250; Santa Cruz), anti-PAX6 (mouse, 1:250, Santa Cruz), anti-RELN (mouse, 1:300; Millipore), anti-NeuN (Rabbit,1:1000, Abcam), anti-BrDU (mouse, 1:250, BD Biosciences), anti-BrDU (rat, 1:250, Accurate Chemicals). For the quantitative analysis of total number of BCL11B+ and SATB2+ cells, z-stack images from (n=3/ species) were utilized to segment DAPI signals from individual nuclei, and fluorescence signals from the nuclei were obtained using Volocity (v.6.3.1). The positive signals for each marker protein were defined based on the median nuclear intensity from all cells analyzed by (v.11.2.0) Spotfire software.

Immunoblot analyses were performed using a previously described protocol [73]. Briefly, neocortical tissue from PD 0 *Zbtb18*$^{fl/fl}$; *Emx1-Cre* mice (n=3), *Zbtb18*$^{fl/+}$; *Emx1-Cre* mice (n=3), and wild-type mice (n=3) were isolated and snap frozen in liquid nitrogen. Samples were then minced and mixed with lysis buffer (150 mM NaCl, 1.0% NP-40, 50 mM Tris-HCl pH 8.0) including protease inhibitors (Roche). Equal amounts of protein from each sample were loaded for SDS-PAGE, followed by western blot analysis with signals detected using the ECL detection reagent. Antibody dilutions used for immuno-blot were as following: anti-GAPDH (rabbit, 1:5000; Abcam, ab9485), anti-CUX2 (rabbit, 1:1000, Abcam), anti-ZBTB18 (rabbit, 1:2000, Protein Tech), anti-ZBTB18 (goat, 1:2000, Santa Cruz). For detailed antibody information please refer to Extended Data Table 14.

**Obtaining multi-species alignments of TF binding sites**

Beginning with the lists of putative TF binding sites identified by FIMO as described earlier[66], we selected only those motifs that are recognized by TFs expressed (RPKM>=1) in our RNA-Seq samples derived from either IT or ET neurons for further analysis. This produced a list of 267 motifs. The total number of binding sites for those motifs ranged from 0 to 244,332.

For each one of the 267 motifs, we obtained the multiple-species sequence alignment of sixty vertebrate species from the MULTIZ60 [74] track in the UCSC browser using *mafsInRegion* [65] from UCSC tools. We stitched *maf* alignments and converted them to *fasta* using the script *maf_to_concat_fasta.py* from bx-python (https://github.com/bxlab/bx-python/blob/main/scripts/maf_to_concat_fasta.py).

**Conservation of TF binding sites across species**

We used an in-house script to analyze the multiple sequence alignment of each binding site. First, we calculated the information content for each position in each meme motif using the R package TF binding sites Tools [75]. For each motif, we identified the positions with high information content (>= 0.75) and considered them 'core' positions. Second, we divided the species in the 60-way multiple sequence alignment into four groups: 'Placental mammals, 'Marsupials', 'Monotremes' and 'Non-mammals'. We finally counted, for each binding site, in how many species of each group the motif was present with 0 single nucleotide substitutions in any of the core positions compared to the mouse reference sequence mm10. We repeated the analysis with 1, 2 and 3 mutations. A motif was not considered conserved if it contained an *indel* spanning the core positions.

For each group of species, we produced a binary vector in which, for each binding site, we introduced a 1 if the binding site was conserved and 0 if non-conserved. For placental mammals, we required at least six species fulfilling the conservation criteria stated above. For marsupials, monotremes, and non-mammal species we required at least one species.

**Enrichment in TF binding sites in sets of H3K27ac peaks**

We selected seven sets of ChIP-Seq peaks: peaks annotated to differentially expressed genes between IT and ET neurons, DESeq2 PAdj<0.05 (one set for up and one set for down-regulated genes); peaks differentially enriched between IT and ET neurons as determined by DESeq2 with FC>2 and PAdj<0.1 (one set for IT neuron and one set for ET neuron-biased peaks); peaks annotated to genes differentially expressed between the *Zbtb18* KO or *Zbtb18* heterozygous mice, DESeq2 PAdj<0.05 (one set for KO and one for heterozygous-biased genes); a final set composed by a manually curated list of genes important for CC formation extracted from the literature. In addition, we produced a list of background peaks composed of all H3K27ac peaks which did not fall in any of the previous categories.

For each group of species, we then counted how many conserved and non-conserved binding sites were observed in each set of peaks. We compared the proportion of conserved versus non-conserved binding site to the same ratio in the background peaks by means of a Fisher's exact test. Motifs with FDR-adjusted P-values <0.05 were considered significant.

**Conservation of H3K27ac and ZBTB18 peaks**

We used *mafsInRegion* [65] from UCSC tools to obtain MULTIZ60 alignments for each selected enhancer. We stitched *maf*alignments and converted them to *fasta* using the script *maf_to_concat_fasta.py* from bx-python (bx-python). Pairwise alignment distance between species of each of the selected H3K27ac peaks were obtained using the function dist. alignment from the seqinr package in R.

The intersection of chicken and mouse H3K27ac and ZBTB18 ChIP-seq peaks was conducted using the 'IntersectBed' function within Bedtools. To facilitate this intersection, chicken peaks were initially converted into mouse coordinates (from gg6 to mm10) using LiftOver with -minMatch=0.1. Orthologous regions in chicken of all Zbtb18 ChIP-seq peaks identified in mice were determined using liftover from mm10 to gg6, employing the same minMatch value.

**Cell counting and data analysis**

For each cell counting analysis, neocortical somatosensory regions from three different brains of each condition were used. For PD 0 mouse and late-mid-fetal human brain tissues, 200 μm-wide neocortical columns covering layer I to subplate were used as standard fields to conduct counting. For PCD 14.5 and PCD 16.5 mouse tissues, 200 μm-wide neocortical columns covering layer 1 to ventricular zone (VZ) were used as standard fields. Unpaired two-tailed t-tests were used to detect differences between samples. P<0.05 (Alpha =0.05) was set as the cut-off for significance.

**Replicates**

For ChIP-Seq, 2 replicates were used for each condition. For RNA-Seq, a minimum of 3 replicates were used for each condition. For ChIP-qPCR analysis, a minimum of 4 replicates were used for each condition. For RT-PCR analysis, 3 replicates from each time-point were used for each condition. For *Cux2-E1*, *Cux2-E2*, *Cux2-E3* transgenic mice, 3-7 founders from each line were analyzed. For IUE experiments, 3-6 successfully electroporated animals were examined for each analysis. For IDU/CIDU/BrDU *in vivo* labeling, a minimum of 2 litters were used for each experiment, and 3-5 brains from each litter were examined. For ISH and IF experiments using mouse tissue, a minimum of 3 animals were used for each experiment and between 3 and 20 sections from each animal were examined. For ISH and IF experiments using non-mouse mammals, 2-5 sections from 2 individuals were used. For cell culture experiments, cells were isolated from 3 different mouse brains per condition, and 2 replicates from each brain were used for subsequent observations. For Luciferase assays, a minimum of 4 replicates were used for each transfection and analysis.

### Extended Data References


51 Gong, S. *et al.* A gene expression atlas of the central nervous system based on bacterial artificial chromosomes. *Nature* **425**, 917-925 (2003). https://doi.org/10.1038/nature02033

52 Gorski, J. A. *et al.* Cortical excitatory neurons and glia, but not GABAergic neurons, are produced in the Emx1-expressing lineage. *J Neurosci* **22**, 6309-6314 (2002). https://doi.org/20026564

53 Nakamura, T., Colbert, M. C. & Robbins, J. Neural crest cells retain multipotential characteristics in the developing valves and label the cardiac conduction system. *Circ Res* **98**, 1547-1554 (2006). https://doi.org/10.1161/01.RES.0000227505.19472.69

54 Schwab, M. H. *et al.* Neuronal basic helix-loop-helix proteins (NEX and BETA2/Neuro D) regulate terminal granule cell differentiation in the hippocampus. *J Neurosci* **20**, 3714-3724 (2000).

55 Goebbels, S. *et al.* Genetic targeting of principal neurons in neocortex and hippocampus of NEX-Cre mice. *Genesis* **44**, 611-621 (2006). https://doi.org/10.1002/dvg.20256

56 Sestan, N., Artavanis-Tsakonas, S. & Rakic, P. Contact-dependent inhibition of cortical neurite growth mediated by notch signaling. *Science* **286**, 741-746 (1999).

57 Trapnell, C., Pachter, L. & Salzberg, S. L. TopHat: discovering splice junctions with RNA-Seq. *Bioinformatics* **25**, 1105-1111 (2009). https://doi.org/10.1093/bioinformatics/btp120

58 Langmead, B. & Salzberg, S. L. Fast gapped-read alignment with Bowtie 2. *Nat Methods* **9**, 357-359 (2012). https://doi.org/10.1038/nmeth.1923

59 Zhang, Y. *et al.* Model-based analysis of ChIP-Seq (MACS). *Genome Biol* **9**, R137 (2008). https://doi.org/10.1186/gb-2008-9-9-r137

60 Hamburger, V. & Hamilton, H. L. A series of normal stages in the development of the chick embryo. 1951. *Dev Dyn* **195**, 231-272 (1992). https://doi.org/10.1002/aja.1001950404

61 Matys, V. *et al.* TRANSFAC: transcriptional regulation, from patterns to profiles. *Nucleic Acids Res* **31**, 374-378 (2003). https://doi.org/10.1093/nar/gkg108

62 Messeguer, X. *et al.* PROMO: detection of known transcription regulatory elements using species-tailored searches. *Bioinformatics* **18**, 333-334 (2002). https://doi.org/10.1093/bioinformatics/18.2.333

63 Cartharius, K. *et al.* MatInspector and beyond: promoter analysis based on transcription factor binding sites. *Bioinformatics* **21**, 2933-2942 (2005). https://doi.org/10.1093/bioinformatics/bti473

64 Khan, A. *et al.* JASPAR 2018: update of the open-access database of transcription factor binding profiles and its web framework. *Nucleic Acids Res* **46**, D1284 (2018). https://doi.org/10.1093/nar/gkx1188

65 Kent, W. J. *et al.* The human genome browser at UCSC. *Genome Res* **12**, 996-1006 (2002). https://doi.org/10.1101/gr.229102

66 Grant, C. E., Bailey, T. L. & Noble, W. S. FIMO: scanning for occurrences of a given motif. *Bioinformatics* **27**, 1017-1018 (2011). https://doi.org/10.1093/bioinformatics/btr064

67 Mathelier, A. *et al.* JASPAR 2016: a major expansion and update of the open-access database of transcription factor binding profiles. *Nucleic Acids Res* **44**, D110-115 (2016). https://doi.org/10.1093/nar/gkv1176

68 Ghiasvand, N. M. *et al.* Deletion of a remote enhancer near ATOH7 disrupts retinal neurogenesis, causing NCRNA disease. *Nat Neurosci* **14**, 578-586 (2011). https://doi.org/10.1038/nn.2798

69 Kang, H. J. *et al.* Spatio-temporal transcriptome of the human brain. *Nature* **478**, 483-489 (2011). https://doi.org/10.1038/nature10523

70 Guerrier, S. *et al.* The F-BAR domain of srGAP2 induces membrane protrusions required for neuronal migration and morphogenesis. *Cell* **138**, 990-1004 (2009). https://doi.org/10.1016/j.cell.2009.06.047

71 Hashimoto-Torii, K. *et al.* Interaction between Reelin and Notch signaling regulates neuronal migration in the cerebral cortex. *Neuron* **60**, 273-284 (2008). https://doi.org/10.1016/j.neuron.2008.09.026

72 Sousa, A. M. M. *et al.* Molecular and cellular reorganization of neural circuits in the human lineage. *Science* **358**, 1027-1032 (2017). https://doi.org/10.1126/science.aan3456

73 Kwan, K. Y. *et al.* Species-dependent posttranscriptional regulation of NOS1 by FMRP in the developing cerebral cortex. *Cell* **149**, 899-911 (2012). https://doi.org/10.1016/j.cell.2012.02.060

74 Blanchette, M. *et al.* Aligning multiple genomic sequences with the threaded blockset aligner. *Genome Res* **14**, 708-715 (2004). https://doi.org/10.1101/gr.1933104

75 Tan, G. & Lenhard, B. TFBSTools: an R/bioconductor package for transcription factor binding site analysis. *Bioinformatics* **32**, 1555-1556 (2016). https://doi.org/10.1093/bioinformatics/btw024
